# The RNA cargo of plasma-derived extracellular vesicles in mCRPC patients captures cancer cells and tumor microenvironment signals

**DOI:** 10.64898/2026.09.03.747025

**Authors:** Federico Vannuccini, Yari Ciani, Vera Mugoni, Orsetta Quaini, Francesco Orlando, Veronica Weber, Ariadna Camacho Martinez, Angela Martinelli, Elisa Biada, Alessia Marinelli, Irene Casanova-Salas, Fabio Pessina, Dolores Di Vizio, Alessandro Vannini, Paolo Cremaschi, Stefano Lise, Caterina Nardella, Ugo De Giorgi, Consuelo Buttigliero, Orazio Caffo, Umberto Basso, Joaquin Mateo, Gerhardt Attard, Francesca Demichelis

## Abstract

Circulating analytes in cancer patients capture tumor-related signals. We profiled matched extracellular vesicle (EV) total RNA and cell-free DNA (cfDNA) from the same plasma aliquots of chemo-naive metastatic castration-resistant prostate cancer (mCRPC) patients treated with Enzalutamide (n=54 patients, n=119 longitudinal samples; NCT06981377) at four Italian clinical centers and interrogated data from >10,000 cancer patients’ and healthy individuals’ samples.

Transcript integrity analysis identified coding and non-coding species with fragmentation patterns varying across RNA biotypes. EV-RNA data deconvolution revealed signal from immune populations, including fractions classified as CD4+ T cells, whose abundance increased with disease progression in plasma EVs and mCRPC tissues. By leveraging tissue data-informed mining, we established a novel prostate cancer-related EV-RNA signature that resulted in an independent predictor of poor prognosis and captured tumor microenvironment-derived signals. Integrating EV-RNA and ctDNA information improved patient stratification for progression-free survival. These findings suggest a multifaceted role for plasma EVs as a source of cancer biomarkers.

**Statement of significance:** EV-RNA provides biologically relevant tumor and immune-derived signals in mCRPC. Integration with cfDNA improves patient stratification, highlighting EVs as a complementary source of translational information for monitoring disease progression and therapeutic response in prostate cancer.

## 1 Introduction

Plasma circulating tumor DNA (ctDNA) is the most-studied liquid biopsy analyte for metastatic castration-resistant prostate cancer (mCRPC), accurately capturing the tumor’s somatic genomic landscape, including copy number variations (CNVs), point mutations, rearrangements, and methylation status^1–4^. When combined with serum PSA and clinical characteristics, ctDNA stratification can improve the accuracy of survival prediction in high-volume metastatic disease^5^. Extracellular vesicles (EVs) are in the early stages of translational research but have the potential to expand the scope of liquid biopsies^6^. EVs are lipid-enclosed particles that carry a variety of bioactive molecules, including nucleic acids, proteins, and metabolites^7^. They are secreted by all cells, including tumor cells, into body fluids, and can reflect the molecular characteristics of their cell of origin^8^. In cancer, EVs are involved in processes such as tumor metastasis^9^, immune modulation, and the promotion of chemoresistance^10,11^. The RNA content of EVs, including both coding and non-coding RNAs, has been shown to influence gene expression in distant cells, potentially affecting tumor progression, thereby underscoring their potential to elucidate tumor behavior and guide treatment decisions. Plasma EVs may capture dynamic functional shifts in both malignant epithelial cells and non-neoplastic compartments within the tumor microenvironment (TME)^9^. Recent spatial profiling studies have highlighted the clinical significance of cellular organization within the TME, defining multicellular *spatial ecotypes* associated with malignant progression and patient prognosis^13^. Whether plasma EV-RNA captures the relative abundance of these conserved spatial ecotypes and microenvironmental niches remains largely unexplored.

In this longitudinal multi-center study (NCT06981377), we isolated plasma cfDNA and small (<200nm) EVs^14^ from Enzalutamide-treated mCRPC patients (n=54; 119 samples) to gain complementary insights into the progression of mCRPC and treatment response. Supported by large-scale analyses of 10,690 in-house and publicly available samples data (**Table S1; Fig. 1A**), we showed that i) EV-RNA fragmentation is a non-random, biotype-dependent process, ii) EVs carry both prostate cancer and TME-related signals aligned with prostate cancer tissue signals, and iii) EV information is distinct from ctDNA and associates with patients’ prognosis. By combining ctDNA and EV-RNA profiling, liquid biopsy bridges tumor genomics and EV biology, capturing distinct prognostic signals from both cancer cells and the TME, offering a comprehensive approach for monitoring disease progression in mCRPC.

**Figure 1.**
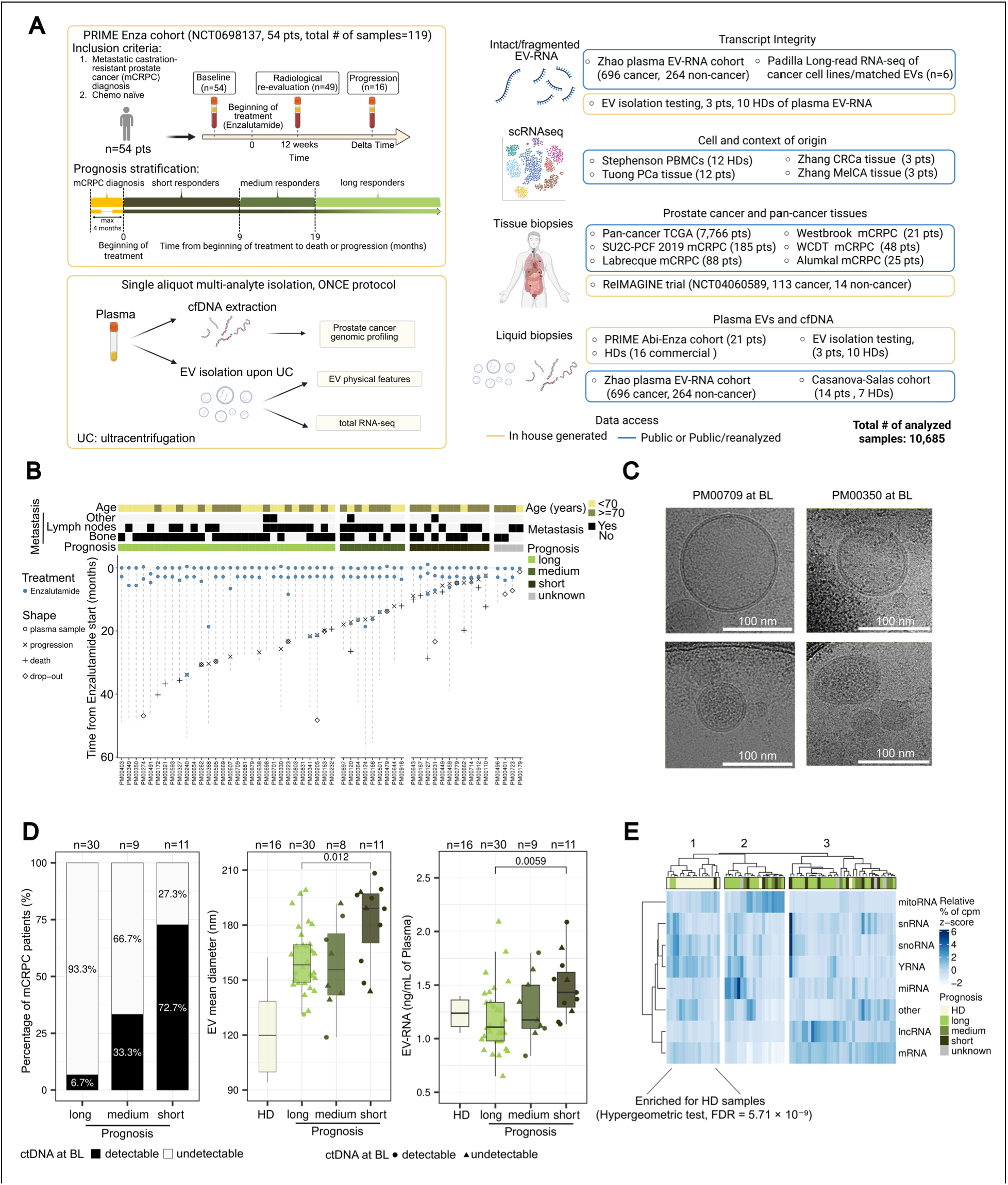
Parallel profiling of plasma EV-RNA and cfDNA of Enzalutamide-Treated Chemo-Naïve mCRPC Patients: A) Top left, sketch of the chemo-naïve mCRPC study cohort (n=54) receiving Enzalutamide as first-line treatment (NCT06981377). Patients are stratified by prognosis as in Fernandez-Perez et al^15^, long responders (n=30), short responders (n=11), and medium responders (n=9). Bottom left, EVs and cfDNA are isolated from the same plasma aliquot using the ONCE protocol^16^. EVs underwent physical and molecular characterization, and the extracted RNA was sequenced via total RNA-seq with a single-end stranded protocol. cfDNA was subjected to a prostate specific targeted NGS assay. Right, summary of total analyzed samples (n=10,685) stratified vertically across analytical domains: transcript integrity, cell and context-of-origin, prostate cancer and pan-cancer tissue, and plasma EVs. Datasets are grouped by data access tier: yellow boundaries indicate in-house data, whereas blue boundaries indicate public or public/reanalyzed data. B) Swimmer plot Enzalutamide-treated mCRPC cohort. The top heatmap displays clinical variables, while the y-axis represents time from treatment initiation. Distinct shapes denote event types (plasma sampling, disease progression, death, and study drop-out). C) Representative cryo-EM images of plasma EVs from two patients at BL, in line with MISEV2023 guidelines14. D) Percentages of patients with detectable ctDNA (left), size of EVs (middle), and EV-RNA quantifications (right) in patients at BL stratified by prognosis. Percentage of patients positive for ctDNA signal (left) is estimated based on genomic alterations using the PCF_SELECT assay and its associated computational pipeline4. Mean diameter size of EVs (middle) is measured using Nanoparticle Tracking Analysis (NTA) on available plasma samples. Plasma-derived EV-RNA has been quantified via Agilent 2100 Bioanalyzer (RNA 6000 Pico Assay). Healthy donors are shown for comparison. Two-sided Wilcoxon Mann-Whitney test p-value is reported. E) Heatmap showing the z-score transformed percentage of cpm for each RNA biotype; hierarchical clustering of the samples is displayed on top. Hypergeometric test for enrichment of HDs in cluster 3 is reported. Abbreviations: mCRPC, metastatic castrationresistant prostate cancer; PCa, prostate cancer; MelCa, melanoma; CRC, colorectal cancer; EV, extracellular vesicle; cfDNA, cell-free DNA; ctDNA, circulating tumor DNA; PFS, progression-free survival; s.d., standard deviation; BL, baseline; HD, healthy donor; NTA, Nanoparticle Tracking Analysis; cpm, counts per million; SE, single-end.

## 2 Results

### 2.1 Multi-analyte characterization of a cohort of first-line mCRPC patients

The observational PRIME study (NCT06981377) recruited patients with advanced prostate cancer across four Italian clinical centers between 2019 and 2026 and collected longitudinal plasma samples, using harmonized protocols. Blood samples were collected from patients at baseline (BL), after 12 weeks of treatment (12w; **Fig. 1A**, top left), and at disease progression (PR) of each therapy. A set of 54 chemotherapy-naïve mCRPC patients who had started Enzalutamide treatment within 4 months of their mCRPC diagnosis (hereafter referred to as the PRIME Enza cohort; **Table 1**) were subjected to targeted cfDNA and total EV-RNA sequencing. Patients were stratified based on progression-free survival (PFS), defined as time to progression or death^15^, as short, medium, and long responders (PFS<9, 9≤PFS<19, and PFS≥19 months, for short, medium, and long, respectively) (**Fig. 1A**, bottom left). Among the PRIME Enza cohort patients, 68.5% had bone metastases, and 54% lymph node metastases (**Table 1**; **Fig. 1B**). An additional set of n=21 mCRPC patients treated with Abiraterone (n=10) or Enzalutamide (n=11) was profiled as part of this study, 16 of which received prior chemotherapy (**Materials and Methods** section **4.2**) (referred to as PRIME Abi-Enza cohort, n=21) together with a control cohort of 16 non-cancer individuals (healthy donors, HDs) for comparative analyses. To comprehensively characterize the plasma EV-RNA landscape of mCRPC patients, we integrated this study dataset with public resources into a broader framework encompassing 10,685 samples, including single-cell RNA sequencing (scRNAseq), cancer tissue cohorts, matched cell and EV transcriptomics, and plasma EVs (**Fig. 1A**, right panel).

cfDNA and EVs were isolated from the same plasma aliquot using the in-house developed ONCE approach^16^ (**Fig. 1A**, bottom left). cfDNA was analyzed through the PCF_SELECT genomic panel^4,5^ for ctDNA estimation, CNVs, and single-nucleotide variants (SNVs) detection. EVs were isolated by ultracentrifugation (UC), and the isolated RNA was analyzed by total RNA sequencing.

In accordance with MISEV2023^14^ and MIBlood-EV^17^ guidelines for the characterization of plasma EVs, we confirmed the presence of small EV proteins (TSG101, Flotillin-1, Flotillin-2, Caveolin-1, CD9, and CD81) in both patients and HDs, as well as the expected co-isolation of contaminants^17^, specifically the albumin protein and the Apolipoprotein A1 (**Fig. S1.1A**; see **Materials and Methods** section **4.1.5**).

Cryo-EM of plasma samples from two representative patients confirmed the presence of intact lipid bilayer-enclosed vesicles with sizes compatible with the expected small EV populations^14^ (<200 nm; as detailed in **Methods 4.3**) (**Fig. 1C**, **Fig. S1.1B**).

Given the established prognostic value of the ctDNA in prostate cancer^1,18^, we tested its association with treatment response. As expected, ctDNA associates with prognosis, with short responders showing a higher frequency of ctDNA positivity (**Fig. 1D**, left) (93.3%, 66.7%, and 27.3% in long, medium, and short responders, respectively). We observed positive correlations between ctDNA, cfDNA concentrations, and ALP at baseline and 12 weeks^19^ (**Fig. S1.2A-B**; consistent with bone metastatic disease in mCRPC.

Baseline EV size (i.e., mean diameter) was larger in short responders compared to long, medium responders, and healthy donors (**Fig. 1D**, middle; **Fig. S1.1C-D**). Notably, short responders also exhibited significantly higher quantities of EV-RNA than other samples (**Fig. 1D**, right). In addition, differences in circulating analytes were observed across time points, with EV diameter decreasing at progression (**Fig. S1.1E**, left) and the number of EVs increasing at the same time point (**Fig. S1.1E**, right). The proportion of patients with detectable ctDNA decreased at 12 weeks and increased again at progression (**Fig. S1.1F**), consistent with an increased tumor burden. Interestingly, no correlation was detected between EV-RNA concentration (**Fig. S1.1G**) and either ctDNA levels or number of EVs (**Fig. S1.2A-C**). Overall, these data suggest that poor responders’ EVs could represent distinct populations that carry a distinct cargo. We therefore sought to comprehensively profile the plasma EV-RNA landscape across the PRIME Enza cohort.

### 2.2 Total RNA landscape of plasma EVs from prostate cancer patients

We sequenced total RNA extracted from plasma-derived EVs of n=135 samples (including longitudinal patient samples and healthy donors). Bioanalyzer profiling of EV-RNAs highlighted the prevalence of short RNA fragments (25 - 200 bp**, Fig. S1.1H**, left, **Table S3**).

We calculated the relative contribution of diverse RNA biotypes and observed that a prominent fraction of the reads mapped to mitochondrial genes, in accordance with recent studies^20^, and to long RNA species such as lncRNAs and mRNAs (**Fig. 1E**). Cluster analyses based on the percentage of counts per million (cpm) associated with RNA biotypes (**Fig. 1E**) strikingly separating healthy donors from mCRPC patients (cluster #3 significantly enriched for healthy donor samples; hypergeometric test, FDR = 5.71 × 10⁻⁹, **Table S4**). Ultimately, these findings suggest that mCRPC is characterized by measurable changes in the diverse RNA species carried by circulating EVs.

### 2.3 EV-RNA fragmentation patterns vary by RNA biotype

Although this and other recent studies^6,21^ revealed the presence of long EV-RNA species, their integrity remains poorly investigated. We therefore assessed plasma EV-RNA integrity in the PRIME Enza cohort employing two complementary *in silico* approaches: *i)* the transcript integrity number (TIN)^22^, based on the assessment of the coverage uniformity along each transcript; *ii)* the proxy of fragment length (PFL), an *ad hoc* metric defined as the observed fragment length relative to the maximum possible length (**Fig. 2A**, left), as detailed in **Supplementary methods 1** and **2**. EV-RNA transcripts with high TIN and high PFL were classified as putatively intact, while those with low TIN and low PFL were interpreted as fragmented EV-RNA molecules (**Fig. 2A**, middle). As different RNA biotypes are known to be selectively packaged into EVs and may exhibit distinct stability profiles^7^, we asked whether EV-RNA integrity differs across RNA classes. Strikingly, our data show evidence that differential fragmentation is associated with RNA biotypes, with YRNA and mitochondrial RNAs being the most intact. In contrast, mRNAs and lncRNAs were highly fragmented (**Fig. 2B, Table S5-6**). Notably, these findings were confirmed in a large, independent, publicly available cohort of plasma EV-RNA samples (n=960), including cancer patients, individuals with benign conditions, and healthy donors^23^ (**Fig. 2C**, **Fig. S2.1A**). Furthermore, plasma EV-RNA fragmentation profiles are independent of EV isolation methods (**Fig. S2.1B, Table S7-8**), and droplet digital PCR (ddPCR) confirmed that transcript abundance trends matched those observed by total RNA-seq (**Fig. S2.1C**, **Table S9**).

**Figure 2:**
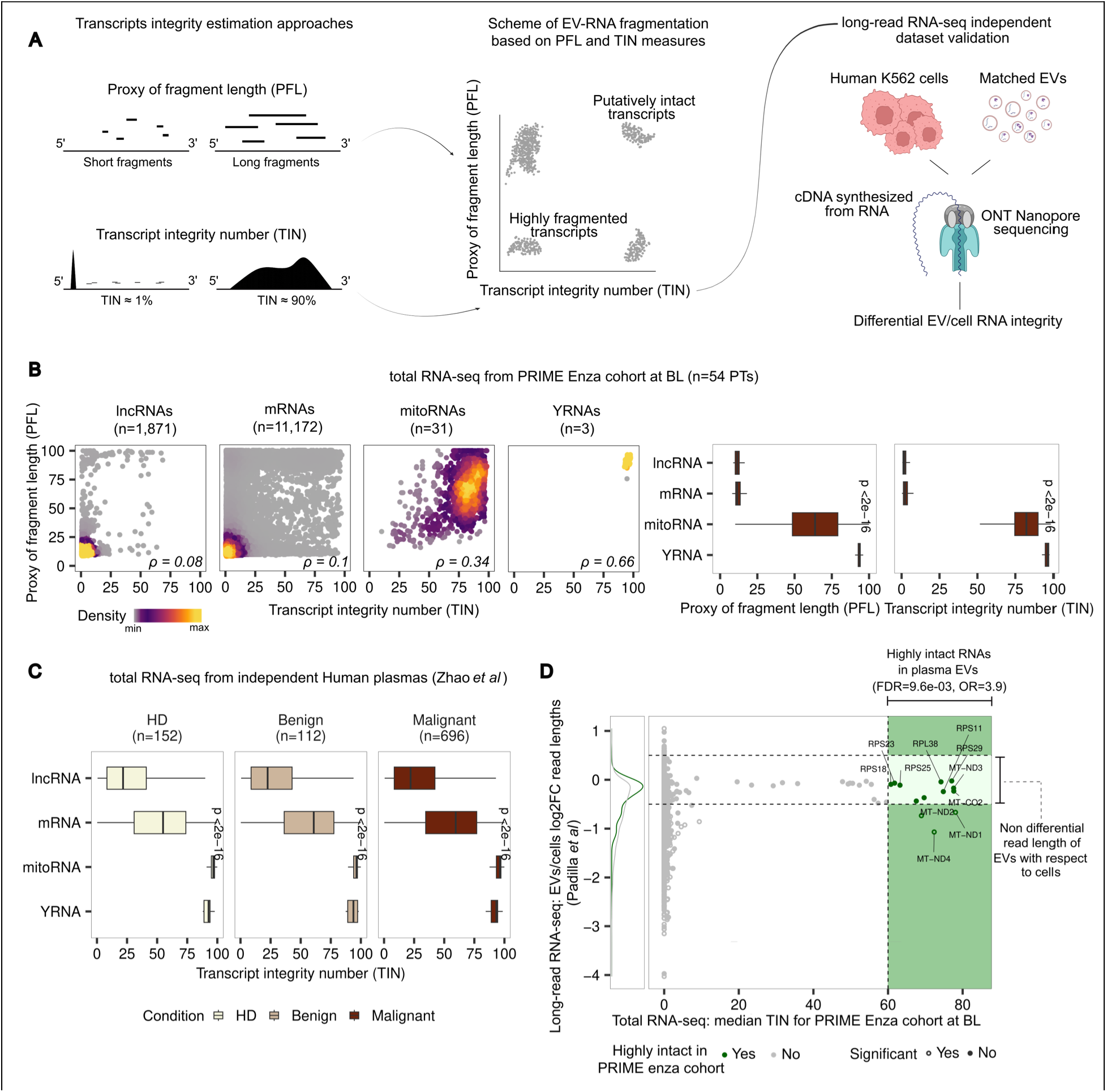
Plasma EV-RNA fragmentation varies by RNA biotype: A) In silico estimation of transcript integrity (left) using two complementary approaches: the Transcript Integrity Number (TIN), based on transcript coverage uniformity^22^ (top), and the Proxy of Fragment Length (PFL), based on read mapping length (bottom). Scheme of EV-RNA fragmentation based on PFL and TIN (middle). Validation using public long-read EV-RNA-seq data from Padilla et al., 2023^24^ (right). B) Integrity measures stratified by RNA biotype. Left, scatter plots of TIN versus PFL with Spearman correlation (gene-sample pairs: lncRNAs = 10,676, mRNAs = 125,787, mitoRNAs = 1,240, YRNAs = 153). Right, boxplots of PFL and TIN distributions across the same biotypes (Kruskal-Wallis test). C) Boxplots of TIN distributions in the independent Zhao plasma EV-RNA cohort^23^ (Kruskal-Wallis test). D) Scatterplot of read lengths log2FC between matched cells and cell-derived EVs from long-read sequencing (y-axis) versus PRIME Enza cohort median TIN at BL (x-axis). Green dots represent transcripts that scored as highly intact in the PRIME Enza cohort, grey otherwise; filled and empty dots represent significant and non-significant differences in read length between EVs and cells, respectively. The green shaded quadrant defines the x-axis threshold for highly intact status. Abbreviations: EV, extracellular vesicle; TIN, Transcript Integrity Number; PFL, Proxy of Fragment Length; lncRNA, long non-coding RNA; mRNA, messenger RNA; mitoRNA, mitochondrial RNA; log2FC, log2 fold change; BL, baseline; Enza, enzalutamide.

To ultimately validate the integrity analysis results, we analyzed a publicly available Nanopore sequencing dataset of cDNA derived from RNA isolated from human K562 cells and matched K562 cell-derived EV-RNA^24^ (**Fig. 2A**, right panel for the Nanopore experimental layout). Nanopore sequencing enables direct assessment of transcript integrity by generating long reads that span full-length cDNA molecules. To this end, we computationally extracted Nanopore-based long-read sequencing data^24^ read lengths for a back-to-back comparative analysis between RNA from K562 cells and their corresponding EV-RNA **(Fig. S2.1E**). Nanopore sequencing confirmed that the high-integrity transcripts detected in our plasma cohort are preserved intact within K562 cell-derived EVs (**Fig. 2D**). To statistically confirm this concordance, the differences in read lengths were evaluated using Fisher’s test across various median TIN stringency thresholds, testing whether high-TIN transcripts were enriched among those with concordant EV–cell read lengths compared to divergent ones (**Fig. S2.1F**, left). Further, the Nanopore data analysis demonstrated that EV-RNA molecules are shorter than the corresponding cellular RNAs (**Fig. S2.1F**, right; **Table S10**), supporting the prominent presence of fragmented transcripts within the EVs isolated from the PRIME Enza cohort’s plasma.

### 2.4 Plasma EV-RNA deconvolution highlights immune populations associated with prognosis

To assess the contributions of different cell types, including blood components, prostate epithelium, and prostate cancer TME to circulating EVs, we performed a deconvolution analysis of plasma EV-RNA, utilizing a reference that leveraged scRNAseq data from prostate cancer tissue^25^ (n=12 samples) and Peripheral Blood Mononuclear Cells (PBMCs)^26^ (n=12 samples, healthy donors) and BayesPrism^27^ (**Fig. 3B**, top) (**Fig. 3A**, **Table S11**). Genes included in the deconvolution analysis were selected as the intersection between the 10,000 most variable genes in the scRNAseq reference and those in the EV-RNA cargo of the PRIME Enza cohort (n=6,064; see **Supplementary methods** section **4.1**). The reference was tested for batch effects across the two integrated scRNAseq datasets, confirming that cell-type clustering was not driven by technical variation (**Fig. S3.1A-B**, see **Supplementary Methods** section **4.2**).

**Figure 3:**
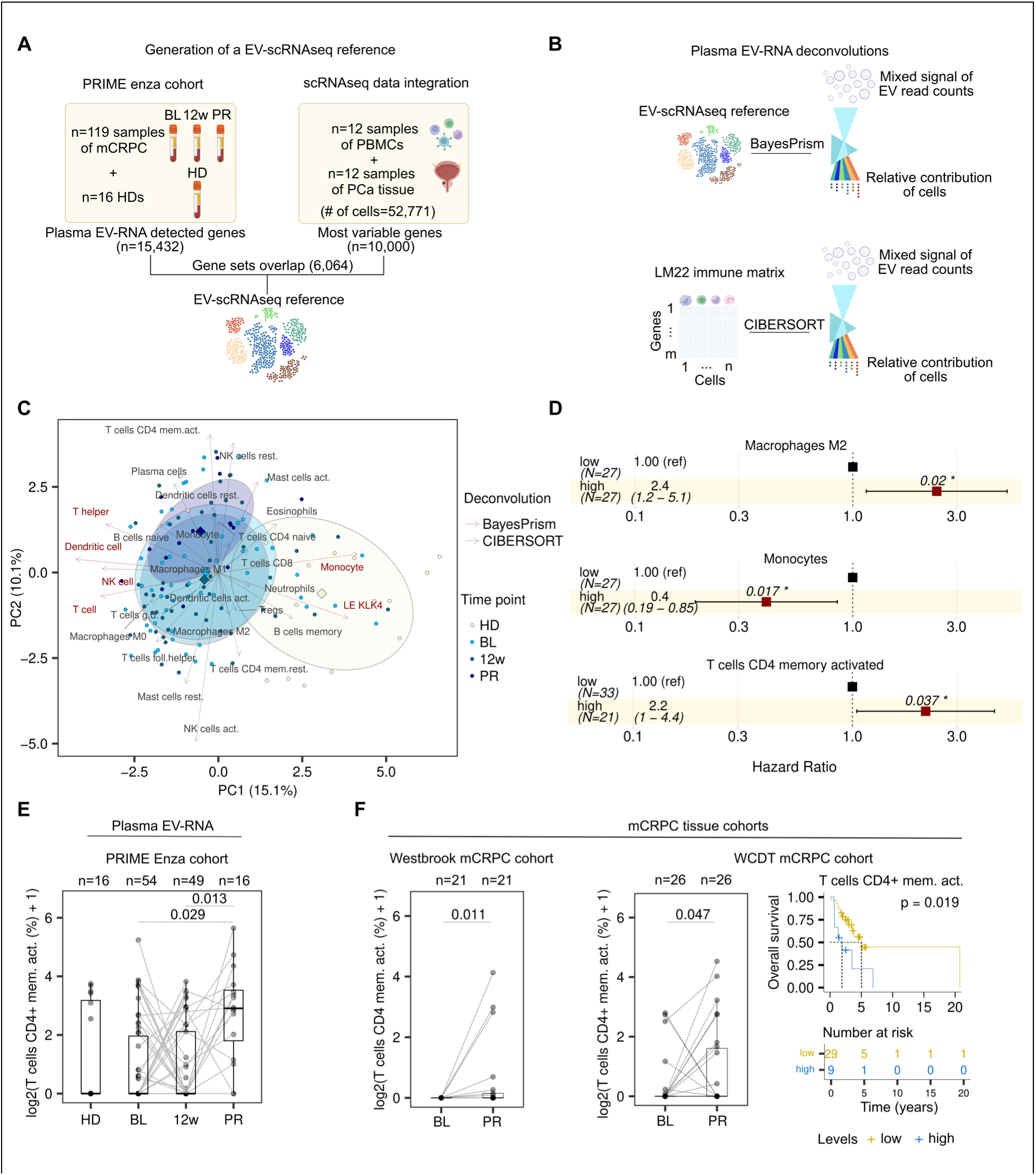
Plasma EV-RNA deconvolution reveals prognostic immune populations, recapitulated in independent datasets: A) Schematic workflow of scRNAseq reference generation using BayesPrism. Genes were first filtered for their representation in the PRIME enza cohort (n=15,432) and then intersected with the top 10,000 highly variable genes identified from an integrated scRNA-seq reference matrix (comprising 52,771 cells from 12 PBMC and 12 PCa tissue samples), nominating a final set of 6,064 genes utilized for plasma EV-RNA deconvolution. B) Top, sketch of plasma EV-RNA deconvolution via BayesPrism using the custom EV-scRNA-seq reference (n=6,064 genes). Bottom, plasma EV-RNA deconvolution via CIBERSORT using the LM22 immune matrix^28^ (n=541 genes). C) Principal component (PC) analysis of plasma EV-RNA deconvolved fractions. BayesPrism and CIBERSORT outputs were concatenated to identify major axes of variation across HDs and patients at different time points. BayesPrism and CIBERSORT loadings are colored in red and gray, respectively. Centroids and ellipses are colored according to HDs status and distinct patient time points. D) Univariate Cox proportional Hazard ratios of statistically significant plasma EV-RNA immune cell fractions deconvolved via CIBERSORT from the PRIME enza cohort. E) CIBERSORT T cells CD4+ memory activated at different time points of PRIME enza cohort and HDs. Two-sided Wilcoxon Mann-Whithney test p-value is reported. F) CIBERSORT T cell CD4+ memory activated fractions in bulk RNA-seq mCRPC tissue cohorts. Left, T cell CD4+ memory activated levels in the Westbrook cohort29 (n=21 pts) at BL and PR, and in the WCDT cohort30 (n=26 pts, including Abiraterone- and Enzalutamide-treated paired patients); paired Wilcoxon’s Mann-Whitney p-values are reported. Right, Kaplan-Meier curve showing overall survival (OS) according to T cell CD4+ memory activated levels in ARSI-naïve patients from the WCDT cohort treated with Enzalutamide (n=38 pts); statistical significance is indicated by log-rank p-value. Abbreviations: EV, extracellular vesicle; scRNA-seq, single-cell RNA sequencing; PBMC, peripheral blood mononuclear cell; PCa, prostate cancer; mCRPC, metastatic castrationresistant prostate cancer; PC, principal component; HD, healthy donor; HR, hazard ratio; CD, cluster of differentiation; BL, baseline; PR, progression; ARSI, androgen receptor signaling inhibitor; OS, overall survival; Enza, enzalutamide; WCDT, West Coast Dream Team.

Clustering analysis of the deconvolution output revealed a cluster significantly enriched in healthy donors (**Table S12**), with higher levels of the erythrocyte component when compared to the other clusters (**Table S13**), and a cluster enriched in long responders (**Table S12**), highlighting their distinct circulating EV-RNA compositions. Notably, short responders exhibit higher T cell fractions than long responders (**Fig. S3.1D**, **Table S14-S16**). The same analysis was conducted on the independent mCRPC patients’ plasma EV-RNA dataset from Casanova-Salas *et al.*^6^, confirming differences between patients and healthy donors (**Figure S3.1E-F**, **Table S17-18**). Given the differences in cell-type representation across response groups, with higher T-cell proportions in short responders and higher endothelial fractions in long responders (**Table S14-S16; Table S19-S20; Fig. S3.1D**), we opted to further investigate the contribution of immune cells to the repertoire of EV-RNA in circulation. To this end, we performed an additional deconvolution analysis following an approach previously used for plasma EV-RNA^31^, specifically focused on the immune component, using CIBERSORT^28^ and a reference matrix of 22 immune cell types (LM22; **Fig 3B**, bottom). The correlation between the outputs of the two deconvolution methods and blood counts and RNA biotypes (**Figure S3.2-3.3**) shows that each method captures unique, non-overlapping aspects, with CIBERSORT leveraging a reference matrix^28^ that inherently provides higher granularity within immune cell subsets. To explore global patterns of variation, we performed principal component analysis on the concatenated deconvolution outputs from BayesPrism and CIBERSORT (**Fig. 3C**). This analysis revealed that BayesPrism T cells were concordant with CIBERSORT gamma delta T cells (i.e., T cells g. d.), whereas CIBERSORT CD4+ memory activated and resting T cells projected in opposite directions. Furthermore, the first principal component clearly separates healthy donors from patients, with Monocytes and luminal epithelial (LE) KLK4 fractions, estimated with BayesPrism, contributing predominantly to healthy donors’ samples. In contrast, the second principal component captures disease evolution, showing a shift in patients at progression relative to other time points and healthy donors, highlighting CIBERSORT CD4+ memory activated T cells as a major driver of this variation. Together, these results suggest that both immune cells and epithelial signals shape the plasma EV-RNA landscape and its evolution during disease progression in response to therapy.

We next investigated the association between CIBERSORT output and patients’ PFS. High Monocyte levels at baseline are associated with better PFS, whereas elevated Macrophage M2 levels are linked to worse PFS in the PRIME Enza cohort of plasma EV-RNA (**Fig. 3D**, **Table S21**). To systematically evaluate the prognostic potential of deconvoluted cell populations, we assessed all identified immune subsets across tissue cohorts from diverse consortia, comprising **777** prostate cancer samples. Across these datasets, the association of Monocytes with PFS is consistently observed in both the SU2C-PCF 2019^32^ **(Fig. S3.4A**) and West Coast Dream Team (WCDT)^30^ mCRPC tissue cohorts (**Fig. S3.4B**), while the prognostic value of Macrophages M2 is further supported in the PRAD-TCGA^33^ (**Fig. S3.4C**) and Alumkal^34^ tissue cohorts (**Fig. S3.4D**). CD4+ memory activated T cells showed high levels associated with PFS at both baseline (**Fig. 3D**, **Table S21**) and at 12 weeks (**Table S22**), highlighting their potential role in patients’ immune response. Since CD4+ memory activated T cells were originally characterized using *in vitro* cell culture systems under controlled polarization and activation conditions, we compared their profile with an alternative reference of sorted blood cells^35^ and found a positive correlation with regulatory T cells (Tregs; **Figure S3.3E**), supporting their potential involvement in immune suppression^36,37^. Strikingly, this population also exhibited higher levels at progression (**Fig. 3E**, one-tailed Wilcoxon test; **Table S23-25**) with respect to baseline, as confirmed by two independent sets of mCRPC RNA-seq tissue data: in the Westbrook^29^ (**Fig. 3F**, left; **Table S26**) and the WCDT^30^ cohorts (**Fig. 4F**, middle; **Table S27**). Notably, this population is associated with poor overall survival (OS) in patients treated with Enzalutamide, who were treatment-naïve at baseline in the WCDT cohort (**Fig. 3F**, right).

**Figure 4:**
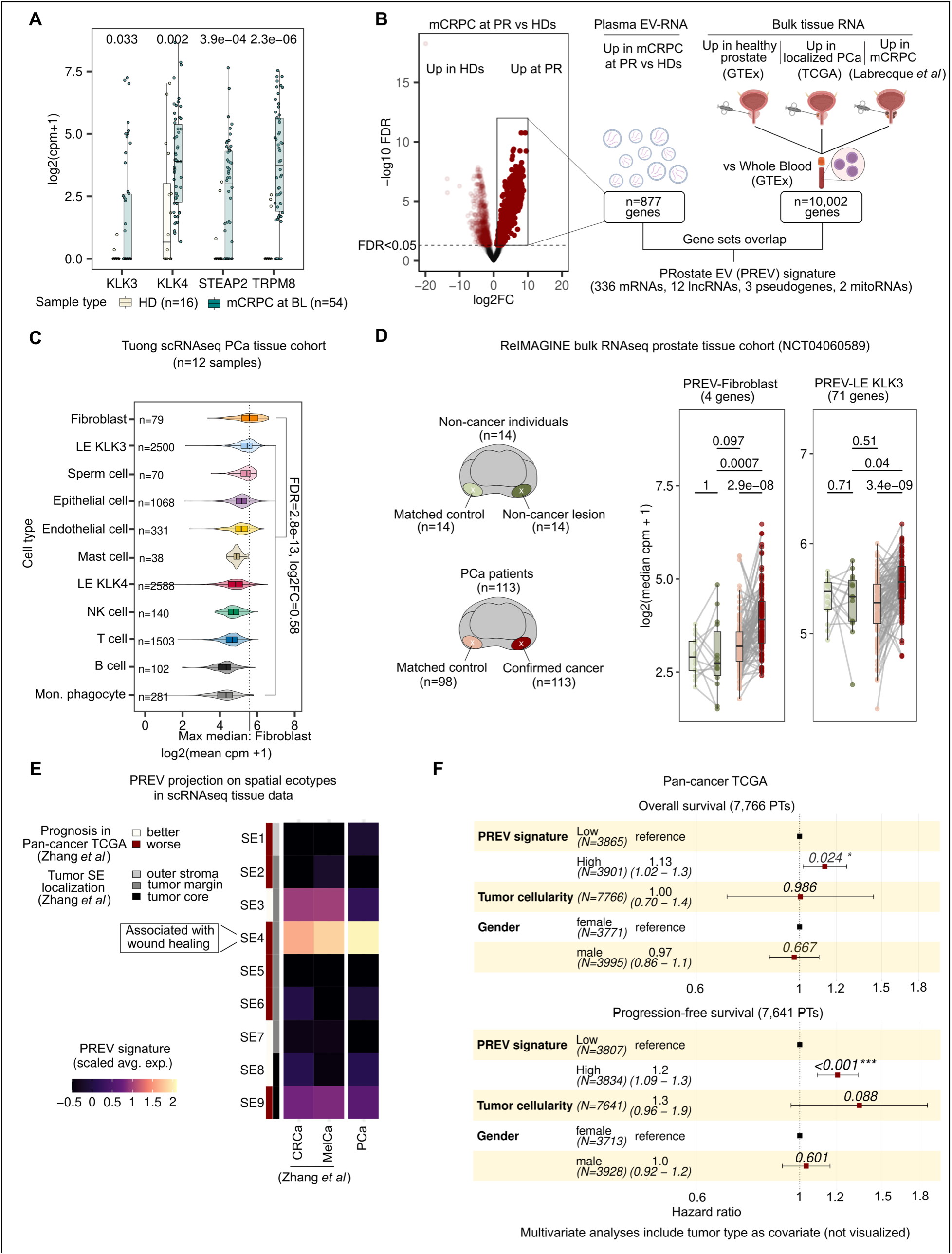
Plasma-derived EVs carry PCa-related signal with prognostic value in tumor tissue: A) PCa-related transcripts are highly represented in plasma EV of mCRPC patients at BL (n=54) with respect to HDs (n=16); Wilcoxon’s Mann-Whitney two sided test is reported on top. B) Left, volcano plot showing genes upregulated in patients at PR (n=877) or in HDs (n=764). Dots are colored by significance (FDR < 0.05). Right, flowchart of PREV signature construction, including 353 genes whose selection is based on differential expression across multiple comparisons: genes upregulated in patients at PR vs HDs (plasma EV-RNA), genes enriched in prostate tissue relative to whole blood (this set results from the intersection of genes upregulated in healthy prostate vs Whole Blood (GTEx^38^), genes upregulated in PRAD-TCGA^33^ vs Whole Blood and genes upregulated in mCRPC^49^ vs Whole Blood). C) Violin plots showing expression of PREV signature across cell types in Tuong scRNAseq PCa tissue cohort^25^; Wilcoxon’s Mann-Whitney two-sided FDR and log2FC is reported on top of violins of fibroblasts with respect to other cells. The maximum median expression of the cell type is reported as a vertical dotted line. D) Left, sketch of the ReIMAGINE prostate tissue cohort (NCT04060589). Right, the expression of cell-type specific PREV genes, selected using the Tuong dataset, including PREV genes highly expressed in Fibroblasts (n=4), LE KLK3 (n=71); Wilcoxon’s Mann-Whitney two-sided paired test is reported for paired individuals, while Mann-Whitney two-sided unpaired test is reported for unpaired biopsies. E) Scaled average PREV signature expression across inferred spatial ecotypes (SEs) from Zhang et al.^13^ in three scRNA-seq datasets, including the Tuong scRNAseq PCa^25^, melanoma (MelCa)^13^, and colorectal cancer (CRCa)^13^; SEs are annotated for tumor localization and Pan-cancer TCGA prognosis, both derived from Zhang et al.^13^. F) Top, multivariate Cox proportional hazard ratio analysis for overall survival of PREV across 27 tumor types included in Ciani et al.^50^, adjusted for tumor type, tumor cellularity, and gender. Bottom, multivariate Cox proportional hazard ratio analysis for PFS of PREV across 26 tumor types included in Ciani et al.^50^, adjusted for tumor type, tumor cellularity, and gender. Abbreviations: PREV, PRostate EV signature; EV, extracellular vesicle; PCa, prostate cancer; mCRPC, metastatic castration-resistant prostate cancer; BL, baseline; HD, healthy donor; PR, progression; FDR, false discovery rate; GTEx, Genotype-Tissue Expression; PRAD, prostate adenocarcinoma; TCGA, The Cancer Genome Atlas; scRNA-seq, single-cell RNA sequencing; log2FC, log2 fold change; LE, luminal epithelial; SEs, spatial ecotypes; MelCa, melanoma; CRCa, colorectal cancer; OS, overall survival; PFS, progression-free survival.

Although correlative, our findings provide consistent and independent evidence across several cohorts supporting the contribution of CD4+ memory activated T cells to the transcriptomic signal observed in plasma EVs from mCRPC patients. In this context, plasma-derived EV-RNA emerges as a valuable source of information on systemic immune modulation.

### 2.5 Plasma EVs carry transcripts associated with prostate cancer

We then sought to identify the fraction of the plasma EV-RNA signal originating from prostate cancer cells in the PRIME Enza cohort. Therefore, we initially focused on genes known for their prostate specificity, relevance to prostate cancer biology, and upregulation in mCRPC patients compared to whole blood healthy tissue from the GTEx database^38^ (**Figure S4.1B**). Specifically, we investigated *KLK3* that encodes PSA, a widely recognized marker for prostate cancer^39^; *KLK4*, which plays roles in androgen receptor (AR) signaling^40^; *STEAP2,* known to contribute to cell proliferation and migration in prostate cancer^41^; and *TRPM8*, whose overexpression induces prostate tumor growth^42^, both previously reported as secreted into cancer cell EVs^43,44^. We first tested their levels in EV-RNA isolated from a panel of androgen receptor (AR) positive and negative prostate cancer cell lines, by ddPCR; as expected, we observed significant abundance in AR-positive cell-derived EVs (i.e., VCaP, LNCaP, and 22Rv1) versus AR-negative ones (**Figure S4.1A**).

Importantly, we noticed that the EV-RNA levels of these selected genes were significantly higher in the plasma samples from PRIME Enza patients compared to those from healthy donors, supporting the presence of cancer-derived signal in plasma EV-RNA (**Fig. 4A**).

Interestingly, EV-RNA sequencing analysis revealed that high levels of *KLK4* and of the AR gene signature^45^ show an association with PFS in patients at baseline (**Fig. S4.1C-D, Table S28**), linking EV-RNA to their putative cell of origin transcriptional program and, in turn, to patients’ prognosis. Notably, when combined with blood ALP and PSA levels (commonly used as diagnostic^46^ and prognostic^47^ biomarkers for prostate cancer^48^, respectively), the prognostic signal of the EV-RNA AR signature is enhanced compared to that of each variable alone **(Fig. S4.1E**, **Table S29**).

To further verify the detection of tumor-derived signals in plasma EVs, we assessed the levels of oncogene transcripts in the presence of genomic DNA amplification in matched cfDNA, profiled using the PCF_SELECT computational pipeline^4,5^ (**Fig. S4.2**). Patients stratified by genomic amplification status at baseline exhibited levels of corresponding oncogene transcripts higher than those of patients with wild-type status and healthy donors (**Fig. S4.1F**). Overall, our findings support the potential of patients’ plasma EV-RNA content in providing tumor-specific information.

### 2.6 Definition of an EV-RNA gene signature to capture tumor-related signal in the circulation

Based on the evidence that EV-RNA carries tumor-specific information with prognostic potential, we applied a data-driven approach to define a prostate cancer-associated plasma EV signature. As an increased percentage of patients exhibited ctDNA positivity at progression compared to baseline and 12 weeks (**Fig. S1.1F**), reflecting enhanced tumor signal in circulation, we performed a differential EV-RNA analysis comparing the high-burden state at progression against healthy donors to capture the tumor-specific plasma EV-RNA repertoire. We identified 877 genes that were differentially abundant significantly (**Fig. 4B**, left). To effectively separate the tumor-derived signal from hematopoietic components, we intersected these genes with bulk RNA-seq tissue data to retain only those that showed a prostate tissue-specific rather than whole-blood-related expression profile (**Fig. 4B**, right; n=10,002). Specifically, three gene sets were derived by the intersection of different comparisons, identifying genes differentially expressed between healthy prostate^38^, PRAD-TCGA^33^ and mCRPC tissue^15^ versus whole blood^38^. This resulted in the PRostate EV signature (PREV; **Table S30**); a set of 353 genes, comprising 336 mRNAs, 12 lncRNAs, 3 pseudogenes, and 2 mitoRNAs. To determine the cellular origin, we investigated the levels of PREV expression in a prostate cancer tissue scRNAseq dataset^25^. We observed high expression in fibroblasts and luminal epithelial KLK3 (LE KLK3) cells (**Fig. 4C**; **Tables S31-S32**), suggesting that transcripts included in the PREV signature may derive from both the TME and the prostate cancer epithelium. Motivated by this multicellular origin, we used the same scRNA-seq dataset to dissect the PREV signature into cell-type-specific gene modules (**Table S33**). Strikingly, in the ReIMAGINE cohort (NCT04060589), a prospective study of men undergoing diagnostic biopsy for suspected prostate cancer (**Fig. 4D**, left), the PREV-Fibroblast and PREV-LE KLK3 modules (**Fig. 4D**, right; **Table S34**) were significantly upregulated in the confirmed prostate cancer biopsies compared to their matched controls. Furthermore, expression of these modules in confirmed tumors was significantly higher than in both the non-cancer lesion and matched control tissues from non-cancer individuals. In contrast, the PREV-LE KLK4 (**Fig. S5.1A**), NK cell, T cell, mast cell, and endothelial modules exhibited an opposite trend, showing reduced expression in malignant tissues (**Table S34**). Collectively, these results suggest that PREV captures a cancer-driven signal originating from both the fibroblasts and the malignant epithelium in prostate cancer patients. Furthermore, the PREV is robustly detectable in epithelial CTCs from Sharifi *et al.*^51^, with its expression significantly enriched in high-purity samples compared to low-purity counterparts (**Fig. S5.1B**). Given that malignant tissues are complex cellular ecosystems rather than isolated cell populations, we next sought to determine the precise origins of the PREV signature within the spatial context of the tumor microenvironment. Interrogation of scRNA-seq datasets spanning prostate cancer^25^, melanoma (MelCa)^13^, and colorectal cancer (CRCa)^13^ revealed a striking enrichment of PREV expression within a specific multicellular program, localized between tumor and stroma, namely spatial ecotype 4 (SE4) (**Fig. 4E**). As previously delineated by Zhang *et al*^13^, SE4 is characterized by the co-occurrence of myofibroblasts and hypoxia-associated endothelial cells, representing an aggressive stromal-endothelial niche associated pan-cancer with poor prognosis and resistance to immunotherapy^13^. Prompted by this clear link to an unfavorable spatial phenotype, we evaluated the broader clinical relevance of the PREV signature using bulk tissue RNA-seq data from the pan-cancer TCGA atlas. Validating the SE4-derived biological rationale, high PREV expression significantly associates with both worse OS (**Fig. 4F**, top; **Table S35**) and PFS (**Fig. 4F**, bottom; **Table S36**) across the pan-cancer cohort. Collectively, these findings demonstrate that the PREV signature captures a highly conserved, tumor microenvironment-associated transcriptional program that is shared across distinct malignancies and tightly linked to more aggressive disease biology.

### 2.7 The clinical relevance of the EV-RNA gene signature in mCRPC patients’ plasmas

Strikingly, we observed high levels of the signature associated with poorer PFS in patients with undetectable ctDNA (ctDNA-/high, **Fig. 5A**), suggesting that plasma EV-RNA may provide valuable prognostic information in mCRPC by capturing tumor-related signals that might otherwise remain undetected when relying solely on ctDNA. Notably, when combined with blood ALP and lymphocytes levels in the same patients’ plasma, the prognostic value of the PREV signature is enhanced compared to these variables alone (**Fig. S5.1C**, **Table S37**).

**Figure 5.**
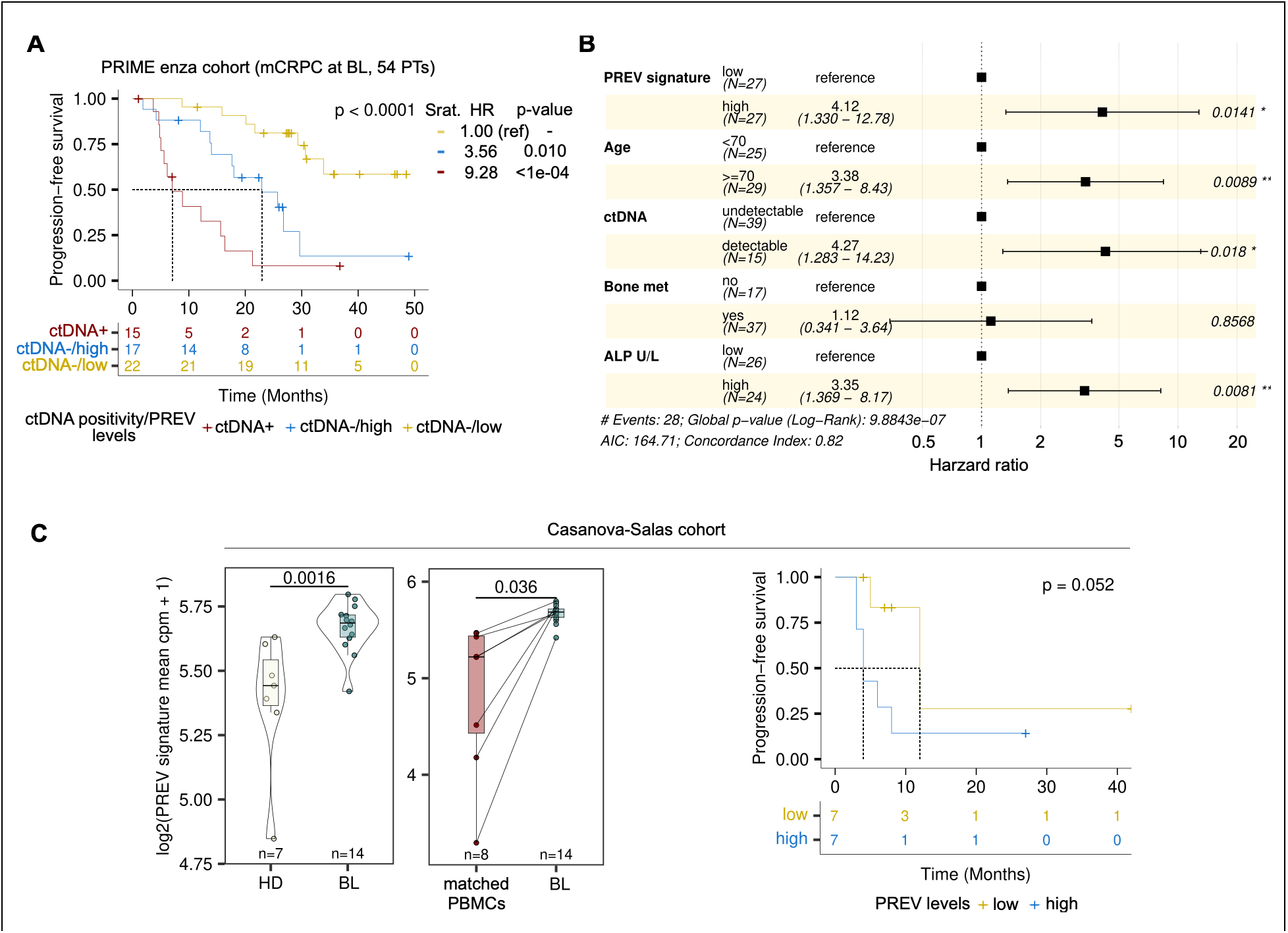
The prognostic value of the PREV signature in mCRPC plasma EV-RNA cohorts: A) Kaplan-Meier curves of PREV levels (n=54) and ctDNA positivity at BL in PRIME Enza cohort. Statistical significance is reported above the curves as log-rank p-value, while univariate Cox proportional Hazard ratios and the corresponding p-values are reported on the right. B) Multivariate Cox proportional Hazard ratio analysis in the PRIME Enza cohort at BL (n=54). Clinical variables have been included based on significance in univariate analysis. C) Left, boxplots representing PREV signature levels at BL in patients (n=14) compared to HDs (n = 7), and relative to matched PBMCs (n=8) within the Casanova-Salas cohort6. Unpaired and paired Wilcoxon Mann Whitney tests were used for patients vs HDs and patients vs matched PBMC comparisons, respectively. Right, Kaplan-Meier curves stratified by PREV levels at BL (n = 14). Statistical significance was determined by log-rank test. Abbreviations: PREV, PRostate EV signature; mCRPC, metastatic castration-resistant prostate cancer; EV, extracellular vesicle; ctDNA, circulating tumor DNA; BL, baseline; HR, hazard ratio; Enza, enzalutamide; HD, healthy donor; PBMC, peripheral blood mononuclear cell.

Unsupervised clustering based on the PREV signature stratified patients in the SU2C-PCF 2019 taxane-naïve mCRPC tissue cohort^32^ into clusters significantly enriched for specific sites of metastasis (**Fig. S5.3A-C**); although not observed in the PRIME Enza cohort (**Fig. S5.2A-B**). Strikingly, multivariate Cox regression identified the PREV signature, age, ctDNA, and ALP as independent prognostic factors in the PRIME Enza cohort (**Fig. 5B**, **Table S38**), suggesting that plasma EV-RNA captures a distinct dimension of tumor progression, maintaining its predictive value independent of standard clinical markers^46^ and ctDNA^1,52^. This prognostic trend was maintained in an additional cohort, namely the PRIME Abi-Enza cohort (n=21; **Fig. S5.1D**, **Table S39**), which is heterogeneous in terms of treatment regimens (Enzalutamide/Abiraterone), with the majority of patients having received prior chemotherapy (n=16; see **Materials and Methods** section **4.2**). Notably, the PREV signature showed no significant difference by ctDNA positivity status in the PRIME Enza cohort (**Fig. S5.1E**, left) and remained distinctly elevated compared to HDs across all time points (**Fig. S5.1E**, right).

Tested in an independent mCRPC plasma EV-RNA dataset^6^ of prostate cancer patients selected for high plasma ctDNA levels, the PREV signature showed significantly higher levels in mCRPC patients at baseline compared to the original study healthy donors and peripheral blood mononuclear cells (PBMCs) isolated from the same plasma samples (**Fig. 5C**, left); its level is elevated at 14 weeks (14w) and at progression with respect to healthy donors and PBMCs (**Fig. S5.1F**), and high PREV levels were associated with poorer PFS in this independent cohort (**Fig. 5C**, right). Notably, despite the high ctDNA burden features, in contrast with the full spectrum of ctDNA levels in our study (**Fig. 1E**), the EV-RNA signature consistently predicted progression, suggesting that it captures a robust, tumor-related signal regardless of the ctDNA content in circulation.

## 3 Discussion

In this study, we generate evidence that integrating plasma EV-RNA transcriptomics with cfDNA genomic profiling enhances risk stratification in chemo-naïve mCRPC patients. Multi-modal liquid biopsy strategies have primarily leveraged multiple analytes as additive tools to maximize sensitivity for tumor-intrinsic signals^6,16^. Here, we extend this framework by demonstrating that EV-RNA and cfDNA capture distinct, orthogonal biological axes. While cfDNA tracks somatic genomic alterations and clonal evolution^5,53–55^, EV-RNA provides a dynamic readout of functional cell states, immune cells, and TME interactions.

This is particularly intriguing when considering that cfDNA might be uninformative for a fraction of patients (93.3% at baseline and 27.3% at progression in our study, **Fig. 1E**, left), thus opening the possibility of EV-based reclassification of patients negative for ctDNA, leveraging both cancer- and TME-derived signals. This insight was enabled by our study cohort selection (NCT06981377), which avoided pre-filtering patients based on ctDNA fraction (as applied in previous studies^6^), thereby capturing the full spectrum of ctDNA shedding in a highly curated, clinically homogenous mCRPC cohort.

Unlocking this comprehensive biological readout required moving beyond standard EV transcriptomic workflows. Total RNA-seq analysis of the PRIME Enza cohort detected both coding and non-coding RNAs, providing a more comprehensive view of the EV transcriptomic landscape that expands upon recent studies on plasma EV-RNA^56^. This adds to previous work on mCRPC that relied solely on poly-A-enriched sequencing, thereby excluding non-coding RNAs that may lack poly-A tails and play important regulatory roles.

Within the EV cargo, we observed a remarkable presence of long species in fragmented form. This finding was confirmed by analysis of the Padilla dataset profiled using Nanopore long-read sequencing technology^24^, overcoming the limitations of our experimental design (SE150 protocol). Additionally, we showed that transcript integrity depends on RNA biotype, suggesting that the observed fragmentation is not a random process but is likely regulated by other molecular machinery, for instance, RNA-binding proteins (RBPs) ^58^and RNA modifications. While polyA-based methods are effective for detecting biologically and clinically relevant signals in EV-RNA^6^, including intact molecules^60^, in this study we show that a comprehensive analysis of the EV-transcriptome requires the inclusion of various RNA species, especially fragmented ones.

Plasma EV-RNA deconvolution revealed a systemic shift in mCRPC, transitioning from a homeostatic erythroid baseline to tumor-derived epithelial (*KLK3+* cells) and immune-enriched signals, as confirmed by our independent reanalysis of the Casanova-Salas cohort^6^. Notably, circulating EVs from Enzalutamide short-responders were enriched for CD4+ memory T cell contributions. Because this plasma profile directly mirrors transcriptomic remodeling in independent mCRPC tissue cohorts, it demonstrates that EV-RNA noninvasively captures the microenvironmental immune evasion that underlies primary ARSI resistance.

To translate these discrete cellular signals into a potential prognostic biomarker, we developed the PRostate EV (PREV) signature to capture integrated prostate cancer tumor-stromal remodeling. It integrates epithelial and stromal signals, providing a non-invasive readout of the multicellular TME. The multicellular origin of PREV was confirmed using tissue data from the prospective ReIMAGINE cohort (NCT04060589), in which the PREV-Fibroblast and PREV-LE KLK3 modules were significantly enriched in confirmed prostate cancer diagnostic biopsies compared with matched control and non-cancer tissues.

Spatially, PREV maps to conserved tumor-stromal niches dominated by myofibroblast activity and extracellular matrix remodeling processes known to orchestrate wound healing and stroma-driven immune exclusion across malignancies. The expression profile of this tumoral niche is associated with poor prognosis across TCGA pan-cancer tissue cohorts^13^, suggesting a possible generalization and applicability of EV-RNA profiling approach to solid tumors other than prostate cancer. Given that myofibroblasts are key orchestrators of stromal immunosuppression, fostering immune exclusion through extracellular matrix remodeling and crosstalk with immunosuppressive populations^61^, the ability to detect tumor-stromal niches-associated signals in plasma EVs extends beyond risk stratification. It positions EV-RNA as a promising liquid biopsy readout for identifying stroma-driven immune evasion, carrying direct implications for predicting resistance to immune checkpoint therapies.

Crucially, the PREV signature remains strongly prognostic in ctDNA-negative patients and operates independently of ctDNA burden. By capturing conserved spatial microenvironments from blood, plasma EV-RNA acts as an essential complementary analyte, rescuing clinical risk stratification when ctDNA shedding falls below technical detection limits.

We identified not only specific transcript markers, but also global EV features associated with prognosis. Specifically, when stratifying patients by time to progression, we observed that short responders tend to show larger plasma EVs. As previously reported^62^, large EVs are associated with greater cancer aggressiveness, supporting the existence of distinct EV subpopulations linked to tumor heterogeneity and clinical outcomes. Although our analytical workflow excludes micron-scale vesicles such as Large Oncosomes (>1 μm)^20,63^, the observed associations of EV size, transcript integrity, and deconvoluted cell-of-origin with prognosis strongly suggest that tumor progression is accompanied by systemic shifts in circulating vesicle subpopulations.

Looking forward, combining EV transcriptomic profiling with physical or immuno-affinity enrichment targeting lineage-specific surface markers - such as PSMA for prostate epithelium or CD4/CD45 for immune subsets^64^ - presents a compelling strategy to overcome bulk plasma background, enabling cell-type-specific resolution in EV-based liquid biopsy.

### 3.1 Limitations of the study

While our study provides a comprehensive multi-modal framework for mCRPC, a few limitations should be noted. First, although key findings were cross-validated in independent tissue and liquid cohorts, our primary trial (NCT06981377) and external plasma datasets remain constrained by relatively modest sample sizes. Furthermore, because participating clinical centers in our primary trial were restricted to Italy, broader prospective studies across diverse ancestral populations are required to establish more global generalizability.

Another limitation concerns the definition of immune cell populations, such as macrophages M2 and T cells CD4+ memory activated, used for the deconvolution analysis. These classifications are primarily derived from *in vitro* studies, in which polarization states and functional characteristics are experimentally defined under controlled conditions^28^. Such subpopulations may not fully reflect the complexity of *in vivo* cell states, which are continuously shaped by the tumor microenvironment and systemic signals^36^.

Furthermore, direct deconvolution of spatial ecotypes from liquid biopsy data - using Liquid EcoTyper^13^ - was not feasible due to the use of a targeted cfDNA sequencing approach and the absence of methylation data. Independent of the computational method, deconvolution approaches, and the link between the PREV signature and the spatial SE4 tumor-stromal niche remain associative. Providing direct evidence of the precise cell of origin of circulating EVs represents an intrinsic challenge across the bulk liquid biopsy field, where sequencing outputs reflect an ensemble average of vesicles and particles released from different biological contexts, processes, and cell populations.

Finally, the SE150 sequencing protocol precluded the detection of back-splice junction reads required to identify circular RNAs (circRNAs). Given that circRNAs play well-established regulatory roles in prostate cancer progression and drug resistance^23,65^, paired-end or long-read EV sequencing will be valuable for evaluating their complementary prognostic utility.

In summary, this study demonstrates that plasma EV-RNA and cfDNA represent complementary, orthogonal biological axes in mCRPC precision oncology. By capturing dynamic tumor-microenvironment remodeling, plasma EV-RNA restores prognostic risk stratification in ctDNA-negative patients. Integrating transcriptomic EV profiling alongside genomic liquid biopsies provides a multi-analyte blueprint to non-invasively monitor treatment resistance and guide personalized therapeutic interventions in advanced prostate cancer.

## 4 Materials and Methods

### 4.1 PRIME Enza mCRPC cohort and HDs

#### 4.1.1 Study design and patient enrollment

Metastatic prostate cancer patients were prospectively enrolled in the PRIME observational trial across four Italian clinical institutions; Santa Chiara Hospital (Trento), Istituto Oncologico Veneto (IOV; Padua), Istituto Romagnolo per lo Studio dei Tumori (IRST; Meldola), and Azienda Ospedaliero-Universitaria San Luigi Gonzaga (Orbassano). Chemotherapy-naïve patients who received first-line Enzalutamide within four months of mCRPC diagnosis were selected for the study (n=54; data freeze: 28/02/2024). All patients provided written informed consent. The PRIME Observational Trial (ClinicalTrials.gov ID: NCT06981377) received ethics approval by Santa Chiara Hospital in Trento on 31/01/2019 (Rep. Int. # 2315) and subsequently by the Ethics Committees of all participating centers. Whole blood samples were collected at baseline (n=54), at 12 weeks after Enzalutamide start (n=49), and at disease progression (n=16). Plasma from HDs (n=16, males, with no medical history of cancer or anticancer treatments) purchased from Precision Medicine Group, LLC was included as study control samples. For each patient sample, comprehensive blood count data were collected and stored in the REDCap database (https://projectredcap.org/), including 12 hematologic variables: red blood cells (RBC), hemoglobin (HgB), white blood cells (WBC), platelets, neutrophils, lymphocytes, PSA, testosterone, LDH, ALP and neutrophil to lymphocyte ratio (NLR). Detailed information from longitudinal blood tests, including relevant hematological variables (**Table S2**), has been collected and curated for each patient. An additional 21 patients from the PRIME cohort were investigated, including mCRPC patients treated with Abiraterone (n=10) or Enzalutamide (n=11). All patients treated with Enzalutamide and n=5 treated with Abiraterone had undergone prior chemotherapy. Throughout the manuscript, we refer to this cohort as the PRIME Abi-Enza cohort.

#### 4.1.2 Plasma separation from whole blood collection

All whole blood was collected into K2EDTA-containing tubes (Vacuette, Greiner Bio-One) and stored at 4°C until processing. Plasma from K2EDTA tubes was separated within 2 hours from blood collection by two-step centrifugation at 4°C (1,600 g x 15 min; 3,000 g x 10 min). After isolation, plasma was aliquoted (1.8 mL/vial) and stored at −80°C. The buffy coat was collected after the first centrifuge, divided into 250 μL aliquots, and stored at −80°C.

#### 4.1.3 Concomitant isolation of EVs and cfDNA from human plasma via ONCE approach

After thawing, the plasma samples were filtered (using a 0.8 μm pore) and cleaned from cell debris and larger particles by two serial centrifugation steps: 2,000 g (K-factor 5,340) for 10 min and 10,000 g (K-factor 1,046) for 30 min on a Minispin plus centrifuge (Eppendorf) with F-45-12-11 rotor. EVs were separated and pelleted at 100,000 g (K-factor 122) for 70 min, washed with 1 mL of 1 x PBS (Gibco), and re-pelleted at 100,000 g for 70 min as previously reported^66^ on an Optima MAX-XP ultracentrifuge (Beckman Coulter) equipped with a TLA55 rotor. The liquid fractions (plasma and washing PBS) recovered from the UC steps were pooled in a separate clean tube and stored for subsequent cfDNA extraction according to the ONCE procedure^16^.

#### 4.1.4 Quantification of EVs by Nanoparticle Tracking Analysis (NTA)

Plasma EV size distributions and concentrations were determined for both mCRPC patients and HDs using NanoSight NS300 (Malvern Panalytical). All EV samples were diluted 1:40-1:2,000 in 1x PBS without calcium and magnesium buffer (Gibco) to obtain between 20 and 120 particles/frame. For each sample, 5×60s videos were recorded in standard mode with the equipped sCMOS camera and analyzed by NTA 3.4 software (Malvern Panalytical) with a detection threshold ranging from 3 to 5.

#### 4.1.5 Characterization of EV proteins via Western Blot analysis

Plasma EV proteins were extracted by lysing EVs in 1x RIPA buffer (Millipore) supplemented with protease and phosphatase inhibitors (Roche). Two volumes of lysis buffer were added to one volume of EVs and incubated on ice for 30 min followed by centrifugation at 12,000 g for 15 min at 4°C. Proteins were concentrated by using Amicon ultra centrifugal devices MWCO 10KDa (Merck), and concentration was assessed using BCA assay (Pierce). Total proteins (10 μg) were loaded on 4-15% mini-PROTEAN TGX stain-free gels (Bio-Rad). Gels were activated on a ChemiDoc MP Imaging System (Bio-Rad) and transferred on a trans-blot turbo system (Bio-Rad). Membranes were blocked with blocking solution (Sigma-Aldrich) for 2h at RT and incubated overnight at 4°C with the following primary antibodies: rabbit anti-CD9 (#D801A; Cell Signaling Technology; 1/500); mouse anti-Apolipoprotein A1 (#5F4; Cell Signaling Technology; 1/250); rabbit anti-Albumin (#4929; Cell Signaling Technology; 1/1,000), mouse anti-Tsg-101 (ab83; Abcam; 1/1,000), rabbit anti-Flotillin 1 (#18634; Cell Signaling Technology; 1/1,000), rabbit anti-Flotillin 2 (#3436, Cell Signaling Technology; 1/1,000), rabbit anti-CD81 (#56039; Cell Signaling Technology; 1/500), rabbit polyclonal anti-Caveolin 1 (#PA1-064; Invitrogen). Subsequently, membranes were washed three times with TBS with 0.1% Tween-20 (TBS-T) and incubated 1 h at RT with anti-rabbit or anti-mouse IgG, HRP-linked antibodies (#7074, #7076; Cell Signaling Technology; 1/5,000-1/2,500) in Western Blocker Solution (Sigma-Aldrich). Proteins were visualized on a ChemiDoc Imaging System (Bio-Rad) after incubation of membranes with Amersham™ ECL Select™ Western Blotting Detection Reagent (Cytiva).

#### 4.1.6 Cryo-EM on plasma-derived EVs

Cryo-EM was performed on EVs isolated from BL plasma samples of two mCRPC patients (PM00709 and PM00350). Plasma processing and EV isolation were carried out according to the procedures described in sections **4.1.2** and **4.1.3**, and EVs were resuspended in 20 µL of buffer prior to analysis.

For cryo-EM characterization, lacey carbon EM grids were glow-discharged (60 s, 30 mA) in a GloQube Plus system (Quorum). An aliquot of 2 µL of the EV suspension was applied to the carbon side of the grid, blotted for 6.0 s at 24°C and 100% RH, and plunge-frozen using a Vitrobot Mark IV (Thermo Scientific). Imaging was performed on a Glacios Cryo-Transmission Electron Microscope (Thermo Scientific) operating at 200 kV, equipped with a Falcon 4i Direct Electron Detector (Thermo Scientific) and a post-column Imaging Energy Filter Selectris X (Thermo Scientific). To minimize radiation damage during image acquisition, low-dose mode in EPU software was used. Images were acquired at 130,000× magnification (pixel size 0.955 Å), with a C2 aperture of 50 µm, an objective aperture of 70 µm, and a defocus value of −3 µm. The total accumulated dose per image did not exceed ∼50 e−/Å2. Raw data were processed and analyzed using ImageJ software.

#### 4.1.7 RNA extraction from plasma-derived EVs

RNA was extracted from EVs by qEV-RNA Extraction Kit (IZON Science) with a few modifications to the manufacturer’s protocol. Following EV lysis and column loading, the samples were treated with RNase-free DNase-I (Norgen Biotek) for 5 min at RT. Afterward, the column was washed twice with 600 μL of wash solution. EV-RNA was eluted in 15μL of UltraPure DNase/RNase free distilled water (Invitrogen) pre-heated at 56°C. EV-RNA samples were stored at −80°C. To confirm EV-RNA extraction, the eluted samples were analyzed for distribution of fragment length size and quantified by RNA 6000 Pico Assay (Agilent) on 2100 Bioanalyzer instrument (Agilent) coupled with 2100 Expert version 2.6 software.

#### 4.1.8 cell-free DNA (cfDNA) and genomic DNA (gDNA) extraction

cfDNA was extracted starting from the liquid fractions obtained by the ONCE protocol, as described in paragraph **4.1.3**, using QIAamp Circulating Nucleic Acid kit (QIAGEN) according to the manufacturer’s protocol and eluted in 30 μL Tris HCl 10mM pH 8 (Gibco). The obtained cfDNA was then quantified using Qubit dsDNA High Sensitivity assay (Invitrogen), and the quality was assessed with High Sensitivity DNA assay (Agilent) and analyzed on 2100 Bioanalyzer Instrument (Agilent). gDNA was extracted from 200 μL buffy coat with QIAamp DNA Mini Blood kit (QIAGEN) and eluted in 100 μL Tris HCl 10mM pH 8 (Gibco). The extracted gDNA was quantified using NanoDrop (Thermo Scientific).

### 4.2 EV Isolation methods from prostate cancer patients and HDs plasma, and cell culture medium

Plasma samples were collected and processed according to **4.1.1**, **4.1.2** sections. EVs were isolated through three different methods: UC (according to Thery C. protocol^66^), SEC (qEV/70nm, Izon Science) and CB isolation (according to Notarangelo M. protocol^67^). This selection of patients included 3 prostate cancer patients with replicate samples, along with 10 HDs. Additionally, PC3 cell-derived EVs were included in the study (ATCC, CRL-1435; **Table S7**) cultivated according to the manufacturer’s instructions. Notably, this experiment was DNase-I free, and RNA was extracted by phenol-chloroform precipitation using TRIzol (Invitrogen). The extracted RNA was then purified and concentrated using Single Cell RNA Purification kit (Norgen Biotek Corp.), and quantity and quality were assessed using the RNA 6000 Pico assay (Agilent) on 2100 Bioanalyzer instrument (Agilent) coupled with 2100 Expert version 2.6 software.

### 4.3 cfDNA and gDNA library preparation and sequencing

gDNA for library preparation was fragmented with M220 Focused-ultrasonicator (Covaris) to a target size of 180-220 bp. Libraries for targeted sequencing were prepared starting from 5-25 ng cfDNA and 100 ng gDNA with KAPA HyperPrep Kit (Roche) following the KAPA HyperCap v3.0 protocol with a few modifications. Pre-capture PCR of gDNA was performed for 7 cycles. For probes hybridization, up to 8 cfDNA/gDNA samples were pooled to obtain a combined mass of 2 μg and incubated for capture at 55°C for 16 hours. The captured DNA was then amplified for 10 cycles according to the capture target size. Pre- and post-captured libraries were quantified using Qubit dsDNA High Sensitivity assay (Invitrogen) and the quality was assessed with Bioanalyzer High Sensitivity DNA kit (Agilent). Libraries were then sequenced on a NovaSeq 6000 (Illumina) with a paired-end 150 bp (PE150) protocol.

### 4.4 EV-RNA library preparation and sequencing

VCaP, LNCaP, 22Rv1, PC3, DU145 and RWPE-1 cell lines were obtained from ATCC and cultured according to the manufacturer’s instructions. Each prostate cancer cell line was starved in fetal bovine serum-free medium for 24 hours. After this period, the medium was collected and subjected to sequential centrifugation steps to isolate EVs (300 g for 10 min, 600 g for 5 min, and 2,800 g for 10 min, yielding the 2,8k fraction; 10,000 g for 30 min to obtain the 10k fraction; two rounds of ultracentrifugation at 100,000 × g for 70 min to obtain the 100k fraction). The supernatant was discarded to obtain the small EVs for further analysis. EV-RNA was extracted according to section **4.1.7** (10μL input corresponding to 1-2 ng) and retrotranscribed to random-primed cDNA by using the Reliance Select cDNA Synthesis Kit (Bio-Rad). Freshly generated cDNA (up to 2 μL/reaction) was mixed with 11μL of 2x ddPCR Supermix for probes (No dUTP), 1.1 μL of each target probe (FAM/HEX), and RNAse/DNAse-free water to prepare 22 μL final volume/reaction. Droplets were automatically generated on QX200 AutoDG Droplet Digital PCR System (Bio-Rad) and PCR reaction were run on a C1000 Touch Thermal Cycler (Bio-Rad). A QX200 Droplet Reader (Bio-Rad was utilized for data collection and results were analyzed on QX Manager software (Bio-Rad). Amplicons were generated by using the following corresponding targeted probes: KLK3: dHsaCPE5026548, KLK4: dHSaCPE5027294, STEAP2: dHsaCPE5029993, TRPM8: dHsaCPE5026734, ACTB: dHsaCPE5190200.

Following the same ddPCR setup and workflow established for PCa cell lines, EVs were extracted from healthy donors (HDs). We targeted HBB FTL and GPX1, which were selected as representative plasma whole-blood-specific EV-mRNAs ranking among the top 30 intact transcripts based on TIN in the PRIME Enza cohort at BL (**Table S9**). Amplicons of 81, 68, and 102 bp were generated, respectively, using targeted probe assays from Bio-Rad Laboratories Inc. (HBB: dHsaCPE5044291, FTL: dHsaCPE5046294, GPX1: dHsaCPE5030408). Because successful amplification requires EV-RNA molecules to be at least as long as the targeted amplicon, the detection of these transcripts provides experimental support for the presence of intact EV-RNA molecules.

### 4.6 ReIMAGINE Cohort sample collection and RNA sequencing

Clinical samples were derived from the prospective ReIMAGINE study (NCT04060589), which enrolled treatment-naïve men with suspected prostate cancer based on PSA levels ≤20 ng/mL and a visible lesion on multiparametric magnetic resonance imaging (mpMRI). Alongside standard diagnostic biopsies, participants provided three research cores preserved via the PAXgene Tissue System: two MRI-targeted cores from the index lesion and one spatially distinct non-targeted core from radiologically normal tissue. Central pathological review of hematoxylin and eosin (H&E)-stained sections was performed to evaluate Gleason grade (GG), pattern 4/5 percentage, and tumor cellularity. Cancer cases were selected for RNA sequencing based on diagnostic-research GG concordance (GG ≤2 or GG >2) and research core cellularity >20% (n = 111), alongside non-cancer controls (n = 13). Total RNA (100 ng) underwent rRNA depletion using the NEBNext rRNA Depletion Kit v2 (New England Biolabs), followed by library preparation with the Agilent SureSelect XT HS2 RNA System using 12 PCR cycles (fragmentation was omitted for samples with DV200 < 50%). Pooled equimolar libraries were sequenced on an Illumina platform. RNA-seq was performed on MRI-targeted cores (n = 113 cancer; n = 14 control) and matched non-targeted cores (n = 105 cancer; n = 14 control), excluding samples with insufficient RNA yield or occult tumor detected in non-targeted tissue. All downstream bioinformatic and statistical analyses were conducted on the computing infrastructure at University College of London.

### 4.8 Overview of publicly available cohorts, HDs, and cell lines

#### 4.8.1 scRNA-seq datasets of prostate cancer and PBMCs

PCa scRNA-seq data^25^ were downloaded from https://www.prostatecellatlas.org/# as a single rds file. PBMCs scRNA-seq data^26^ were downloaded from CELLxGENE website: https://cellxgene.cziscience.com/datasets. Spatial ecotypes on prostate cancer data^25^ were inferred as Zhang *et al*^13^, while the corresponding pre-computed ecotypes for MelCa and CRCa were retrieved from the GEO database (GSE320040).

#### 4.8.2 Bulk RNA-seq from healthy tissues and prostate cancer cohorts

Bulk RNA-seq data from male donors across 25 healthy tissues were retrieved from the GTEx^38^ website https://gtexportal.org/home/aboutAdultGtex.

RNA-seq data from prostate cancer tissue samples were obtained from multiple cohorts. The Labrecque cohort^49^ included 88 mCRPC patients without neuroendocrine features, with data downloaded using the recount3 R package. The Westbrook cohort^29^ comprised matched tissue biopsies from 21 mCRPC patients sampled both before Enzalutamide treatment and at progression (42 total samples). The SU2C-PCF 2019 cohort^32^ included 258 mCRPC tissue biopsies from patients either naive (n=163) or exposed (n=95) to AR signaling inhibitors (ARSIs), excluding those treated with taxanes; PFS data were available for 73 patients. For the WCDT mCRPC cohort^30^, the RNA-seq normalized expression matrix from 210 samples was downloaded from https://quigleylab.ucsf.edu/data, and survival data from https://github.com/DavidQuigley/WCDT/tree/master/clinical_metadata.

For localized prostate cancer, the RNA-seq expression matrix from 502 samples in TCGA-PRAD cohort was downloaded via recount3; among these, 501 were localized PRAD and 1 was metastatic. For patients with multiple biopsies, expression values were averaged across biopsies to obtain a single representative profile per patient, resulting in a total of 493 patient-level samples. Patient metadata and genomic estimates were sourced from a previous publication^50^.

#### 4.8.4 EV-RNA-seq datasets

The Zhao cohort of plasma-derived EV-RNA^23^ comprised 152 HDs, 122 individuals with benign conditions, and 696 with malignant disease (across 10 distinct tumor types). The Casanova-Salas mCRPC cohort^6^ included 46 plasma-derived EV-RNA samples collected longitudinally from 14 mCRPC patients treated with ARSI, at baseline (n=14), after 4 weeks (4w) on treatment (n=14), and at progression (n=10), along with PBMCs from 8 patients at BL and plasma from 7 age-matched HDs. Additionally, publicly available EV-RNA data from conditioned media of K562 chronic myelogenous leukemia cells^24^ (n=3 matched EV-cell pairs) were analyzed. These datasets were generated using Nanopore sequencing of polyadenylated full-length transcripts, employing poly-A priming and PCR-cDNA barcoding. EVs were isolated via centrifugation-based filtration followed by SEC.

### 4.9 Plasma-derived EV-RNA-seq data processing

For the PRIME Enza (**4.1**) and PRIME Abi-Enza (**2.2**) cohorts fastq files were trimmed using Cutadapt^68^ v3.5 with parameters suggested from the SMARTer smRNAseq kit instructions: -m 15 -u 3 -a AAAAAAAAAA. Since the samples in **4.1** and **4.2** were sequenced in different batches, the Combat-Seq^69^ v3.35.2 R package was used to correct gene counts for potential batch effect. The resulting matrices were normalized to counts per million (cpm) by using the edgeR^70^ v3.42.4 R package.

For the EV isolation method testing experiment (**4.3**) and the Zhao cohort (**4.7.4**) fastq files were trimmed with fastp^71^ v0.21.0 with parameters: -q 15 -l 15. For all the samples, the reads were mapped to the human genome hg38 (indexed using the GRCh38.primary_assembly.genome.fa and gencode.v41.annotation.gtf files, downloaded from https://www.gencodegenes.org/human/) with STAR v2.7.2b^72^ with parameters: --runThreadN 10 --outFilterScoreMinOverLread 0.66 -- outFilterMatchNminOverLread 0.66 -- chimSegmentMin 15 --chimJunctionOverhangMin 15 -- chimOutJunctionFormat 1 --outSAMtype BAM SortedByCoordinate --quantMode GeneCounts TranscriptomeSAM. The –quantMode tag allowed to generated gene counts, since the software implement HTSeq v2^73^ library.

### 4.10 Estimation of patient ctDNA level and CNVs

Sample ctDNA level and CNVs were estimated as previously described in the original PCF_SELECT manuscript^4,5^.

### 4.11 PRostate EV (PREV) gene signature

The PRostate EV (PREV) signature was developed to identify prostate-specific signal within plasma-derived EVs from mCRPC patients. It includes genes upregulated in patients at PR relative to HDs (n=877), filtered based on prostate tissue-specific expression profile against whole blood (n=10,002) (3 gene sets were derived from comparisons of healthy prostate^38^, PRAD-TCGA^33^ and mCRPC tissue^49^ versus whole blood^38^).

To construct cell-type-specific PREV modules, we leveraged the previously mentioned prostate cancer scRNA-seq dataset^25^ (**4.8.1**). PREV genes were evaluated by comparing each cell type against the rest; those with a log_2_ FC > 1 and FDR < 0.05 were selected. To ensure cell-type specificity, only genes uniquely enriched in a single cell type were retained (**Table S33**)

### 4.12 Plasma EV-RNA deconvolution

Reference-based deconvolution has been performed by using BayesPrism v2.0^27^, leveraging the integrated scRNA-seq data of prostate cancer and PBMCs described in **Supplementary Methods 4** with the parameters: outlier.cut=0.01, outlier.fraction=0.1, Key=NULL. The estimated posterior theta has been used for downstream analysis.

Immune cells reference-based deconvolution has been performed by using CIBERSORT^28^. The R script has been downloaded from https://cibersortx.stanford.edu/ and ran with default parameters. The leukocyte signature matrix (LM22) was used, restricted to the n=541 genes matching those used in the EV-RNA analysis described in section **4.8**.

### 4.13 Statistical analysis

All statistical analyses were conducted using R v4.3.1 on a system running Ubuntu 20.04.06 LTS. For the analysis of sequencing data, raw reads were processed using custom bash scripts. All the plots were done by using the ggplot2 v3.5.1 and ComplexHeatmap v2.16.0 R packages. Statistical tests are indicated in the plots. Correlations were determined using Spearman’s correlation, and the correlation coefficient was used to measure the strength of associations between variables.

For plasma-derived EV-RNA data, differential gene expression analysis was performed by using the DESeq2 v.1.40.2^74^ R package, after accounting for zero-inflation with ZIMBWave and applying log2FC shrinkage using the apeglm algorithm. For the analyses performed on the PRIME Enza cohort, comparisons between patients at progression and HDs, genes with at least 5 cpm in 20% of the samples have been used (**Fig. 4B**, left). For the comparison between short and long responders, genes were included if they met the following criteria: at least 5 cpm in 20% of all samples, along with at least 5 cpm in 70% of samples in short responders, along with at least 5 cpm in 70% of samples in long responders. For the differential gene expression analysis on tissue data (**Fig. 4B**, right), Wilcoxon’s Mann-Whitney two-sided test was used, according to literature when dealing with large sample sizes^75^. All the DEGs in this work have been selected with a filter for |log2FC| >1 and FDR < 0.05.

The Kaplan-Meier analyses, were performed by using survminer v0.4.9.999 and survival v3.5.5 R packages. The log-rank p-values have been computed with respect to time to progression from the start of treatment. All the curves were stratified with respect to the median value of the correspondent variable.

For the PRIME Enza cohort, only variables reaching statistical significance in univariate analysis (p-value < 0.05; **Table S38**) were selected for multivariate Cox regression (**Fig. 5B**). Multivariate Cox regression analyses were performed using the survival v3.5.5 R package, in order to obtain hazard ratios.

## Data Availability

The RNA-seq expression matrices of plasma EV-RNA generated in this study have been deposited on Zenodo (DOI: 10.5281/zenodo.21977463) under restricted access. Raw data will be available on EGA upon peer-reviewed publication.

## Supplementary methods

### 1 Transcripts Integrity Number (TIN)

To assess the integrity of plasma-derived transcripts, the TIN was used as a metric, which evaluates the uniformity of coverage along the length of each transcript^22^. TIN values were computed at the transcript level and subsequently collapsed to the gene level by selecting, for each gene, the transcript with the highest TIN value, thereby retaining the best-supported estimate of transcript integrity.

### 2 Proxy of Fragment Length (PFL)

An additional metric, termed Proxy of Fragment Length (PFL), was used as an orthogonal measure to TIN for assessing transcript integrity in the SE150 sequencing protocol. PFL is based on the read mapping length (RML) from transcriptomic BAM files, with soft-clipped bases excluded when evaluating read length. Specifically, from each file, by using a custom bash script, the CIGAR was extracted and the RML was calculated by using the GenomicAlignments v1.36.0 R package, excluding soft-clipped bases with the *cigarWidthAlongQuerySpace* function. To collapse transcript-level information to the gene-level, PFL is computed as the ratio of the per-gene median RML to the minimum of either the median transcript length from the hg38 annotation or 150 bp (the maximum read length used for the single-end sequencing protocol):

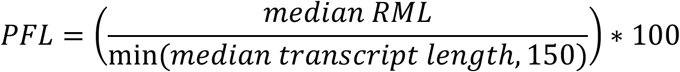

### 3 Nanopore computational workflow for read length analysis

Long-read EV-RNAseq fastq data was downloaded using the fasterq-dump script from SRA toolkit v2.10 using an SRR accession list downloaded from GEO NCBI (https://www.ncbi.nlm.nih.gov/geo/) with accession number GSE225471. Reads adapters were trimmed by using Porechop v0.2.3, then reads were filtered for quality and minimum length by using Chopper v0.8^76^ with the following parameters: -q 7 -l 15. The resulting reads were mapped into the human genome hg38 by using minimap2 v2.28^77^ with parameters: -ax splice. Using the resulting BAM files, the mean and standard deviation of the read lengths were calculated for each sample and used as input for pseudomapping performed with Kallisto v0.44^33^, to assign reads to transcripts, with the following parameters: --single --single-overhang --fr-stranded. To compare the resulting read lengths between cells and EVs, a Wilcoxon’s Mann-Withney two sided test was used.

### 4 scRNA-seq reference contruction for BayesPrism deconvolution

#### 4.1 scRNA-seq data integration

PCa scRNA-seq data^25^ (N = 12) and PBMC scRNA-seq data^26^ (N = 12) were integrated using Seurat v4.3, following published guidelines^78^:

##### Data preparation

The PBMC dataset was streamlined using *DietSeurat*, retaining only count and data slots. The PCa dataset was filtered to include tumor samples only.

Feature selection for integration: *SelectIntegrationFeatures* was used to identify the top 10,000 most variable genes across both datasets, optimizing feature overlap and robustness.

##### Preprocessing

Each dataset was normalized using *NormalizeData* with the Relative Counts (RC) method (scaling factor = 1,000,000). *FindVariableFeatures* (method = "vst") was applied separately to each dataset to select the top 10,000 variable features. Datasets were then scaled and processed using PCA (*RunPCA*). Raw counts were retained for downstream BayesPrism deconvolution.

##### Integration

*FindIntegrationAnchors* and *IntegrateData* were used to integrate the datasets using the shared 10,000 most variable gene set, applying log-normalization.

##### Cell type harmonization

Cell type labels were standardized across datasets (e.g., merging “NK cells” and “nk cell”) to ensure consistency. Harmonized Cell types have been used as Cell Type, while the tissue of origin has been used as cell state.

#### 4.2 Gene filtering based on plasma EV-RNA data

The genes have been filtered to be represented in the PRIME Enza cohort and HDs (see **Materials and Methods** section **4.1**) with 5 cpm in at least 10% of the 135 samples. The 15,432 genes selected were intersected with the 10,000 most variable genes from scRNA-seq integrated data, obtaining 6,064 genes for subsequent deconvolution which has been applied to PRIME Enza cohort.

#### 4.3 Testing for batch effect in scRNA-seq reference

Optimal clustering parameters were determined by sampling random subsets of *n*=5,000 cells from the constructed scRNA-seq reference and applying the shared nearest neighbor (SNN) algorithm at various resolution values. Clustering outcomes were assessed across these resolution values with Silhouette scores, calculated for each cluster to evaluate cohesion. The maximum median Silhouette score allowed to choose the best resolution for clustering: 0.1. The clustering with the optimal resolution was visualized using UMAP, with cells colored by cluster assignment (**Figure S3.1A-B**). Clustering quality was evaluated by calculating the Adjusted Rand Index (ARI) with respect to assay and cell type. A higher ARI with cell type (0.44) than assay (0.15) suggested minimal batch effects related to assay differences, indicating robust clustering by cell type.

## Acknowledgments

The authors would like to thank the CIBIO Department Core Facilities (IRBIO; Next Generation Sequencing and EV biobank), supported by the European Regional Development Fund (ERDF) 2014–2020 and 2021–2027, for their technical support. The authors acknowledge access and services provided by the National Facility for Structural Biology, Fondazione Human Technopole, Milan, Italy. The results published here are in whole or in part based upon data generated by the TCGA Research Network: https://www.cancer.gov/tcga. This work was funded by a Fondazione AIRC per la ricerca sul cancro and Cancer Research UK Accelerator award (22792, to F.D.; A26822, to G.A.); NIH R01CA287075 (to DDV and FD); and by the The John Black Charitable Foundation (to P.C.).

## Notes

https://doi.org/10.5281/zenodo.21977463

### Competing Interest Statement

The authors have declared no competing interest.

