## Supplementary figures for "The RNA cargo of plasma-derived extracellular vesicles in mCRPC patients captures cancer cells and tumor microenvironment signals"

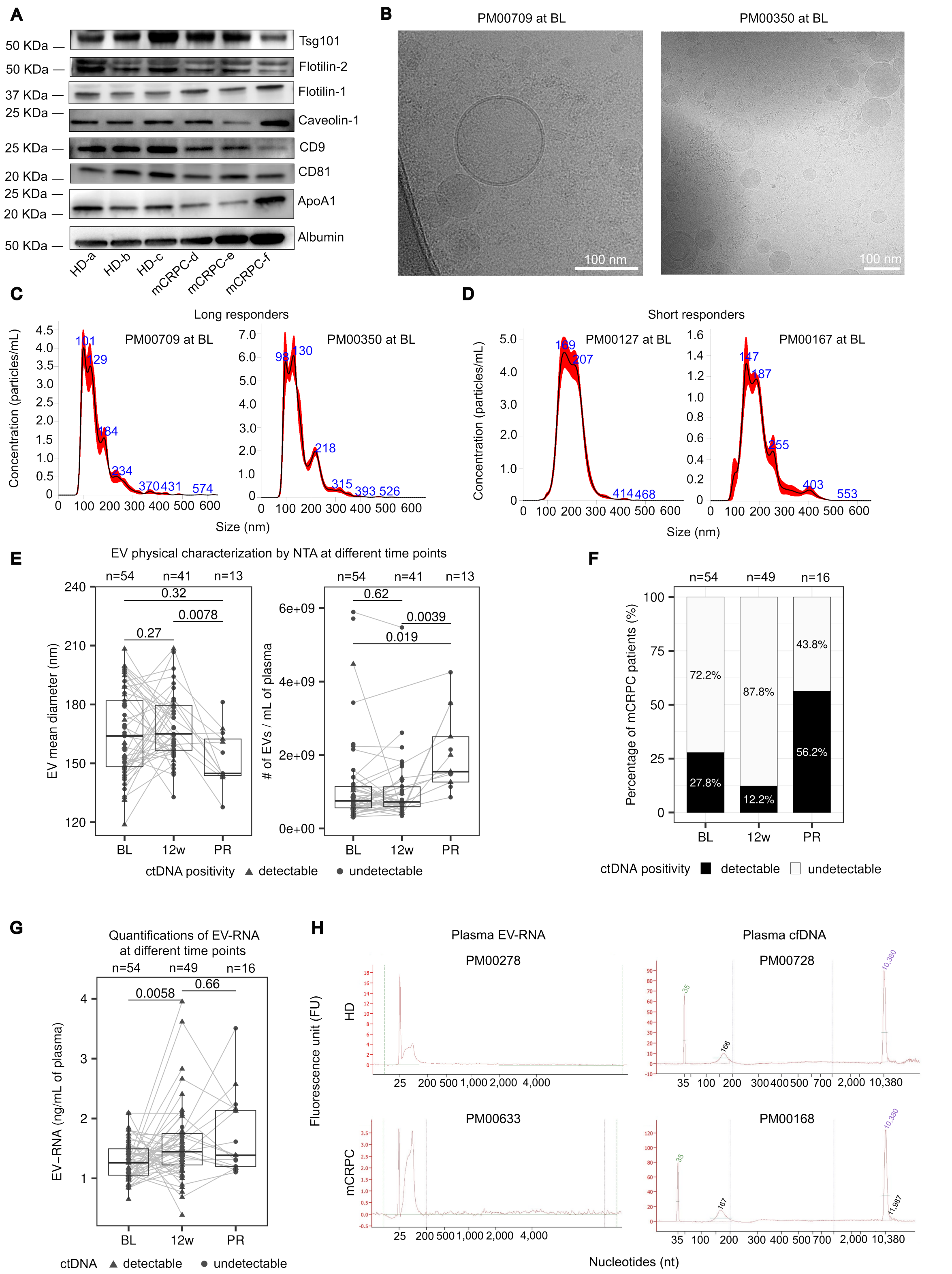


**Supplementary Figure 1.1: Multimodal characterization of EVs at different time points**

A) Representative Western blot according to MISEV2023 guidelines displaying small EV-enriched proteins, including Tsg 101, Flotillins, Tetraspanins (CD9, CD81), and Caveolin 1, along with co-isolated contaminants Apolipoprotein 1 and Albumin. B) Representative uncropped cryo-EM image of plasma-derived EVs from n=2 mCRPC patients. C) Representative EV size distribution from n=2 long responder patients by NTA analysis. D) Representative EV size distribution from n=2 long responder patients by NTA analysis. E) Measurements of the EV size (left) and number of EVs (right) at different time points; two-sided Wilcoxon's paired test is reported. F) Percentages of patients with detectable ctDNA at different time points. Percentage of patients positive for ctDNA signal is estimated based on genomic alternations using the PCF_SELECT assay Statistical enrichment of ctDNA positivity for each time point relative to the remaining cohort was evaluated using two-sided Fisher's exact tests (BL: p-value = 0.67, log2OR = 1.68; 12w: p-value = 0.009, log2OR = -0,40; PR: p-value = 0.004, log2OR = 4.14). G) EV-RNA quantifications at different time points. H) Representative Bioanalyzer profiles of EV-RNA for an HD and a mCRPC patient (left; patient left the study), and cfDNA profiles for an HD and a mCRPC patient (right); two-sided Wilcoxon's paired test is reported. Abbreviations: EV, extracellular vesicle; MISEV, Minimal Information for Studies of Extracellular Vesicles; Tsg101, Tumor susceptibiliy gene 101; CD, cluster of differentiation; cryo-EM, cryo-transmission electron microscopy; mCRPC, metastatic castration-resistant prostate cancer; NTA, Nanoparticle Tracking Analysis; Baseline, BL; 12 weeks, 12w; progression, PR; ctDNA, circulating tumor DNA; cfDNA, cell-free DNA; HD, healthy donor.

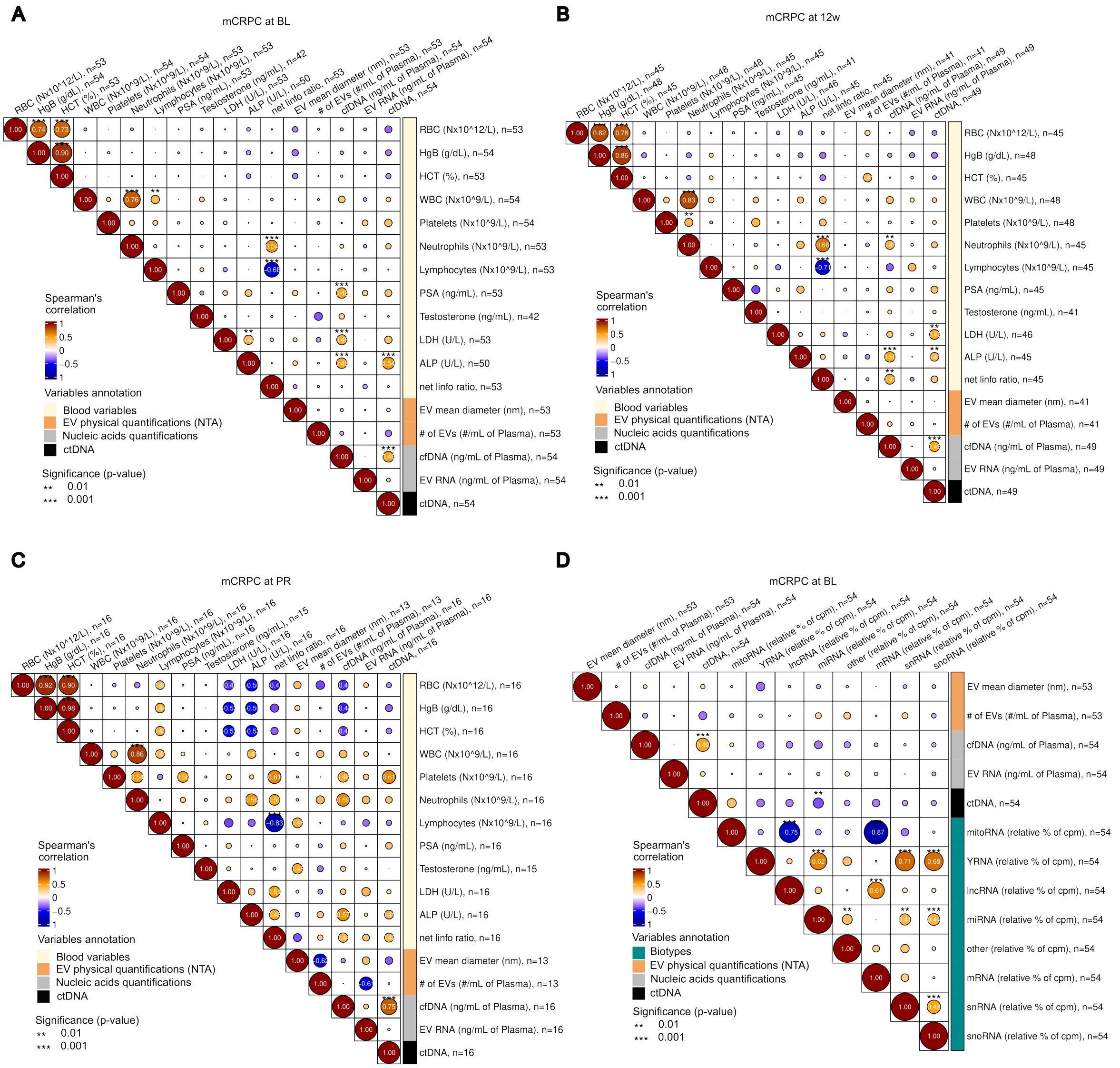


**Supplementary Figure 1.2: Correlation matrices of blood variables, nucleic acids and EV physical parameters**

A) Correlation matrix of blood variables with nucleic acids and EV physical parameters for patients at BL (n=54). B) at 12w (n=49), and C) at PR (n=49). D) Correlation matrix of blood variables and nucleic acids quantifications with respect to the relative % of cpm per biotype. Spearman's correlation test p-values are reported (asterisk). The number of patients for which the observation is measured is indicated in row and column names. Abbreviations**:** EV, extracellular vesicle; BL, baseline; 12w, 12 weeks; PR, progression; cpm, counts per million.

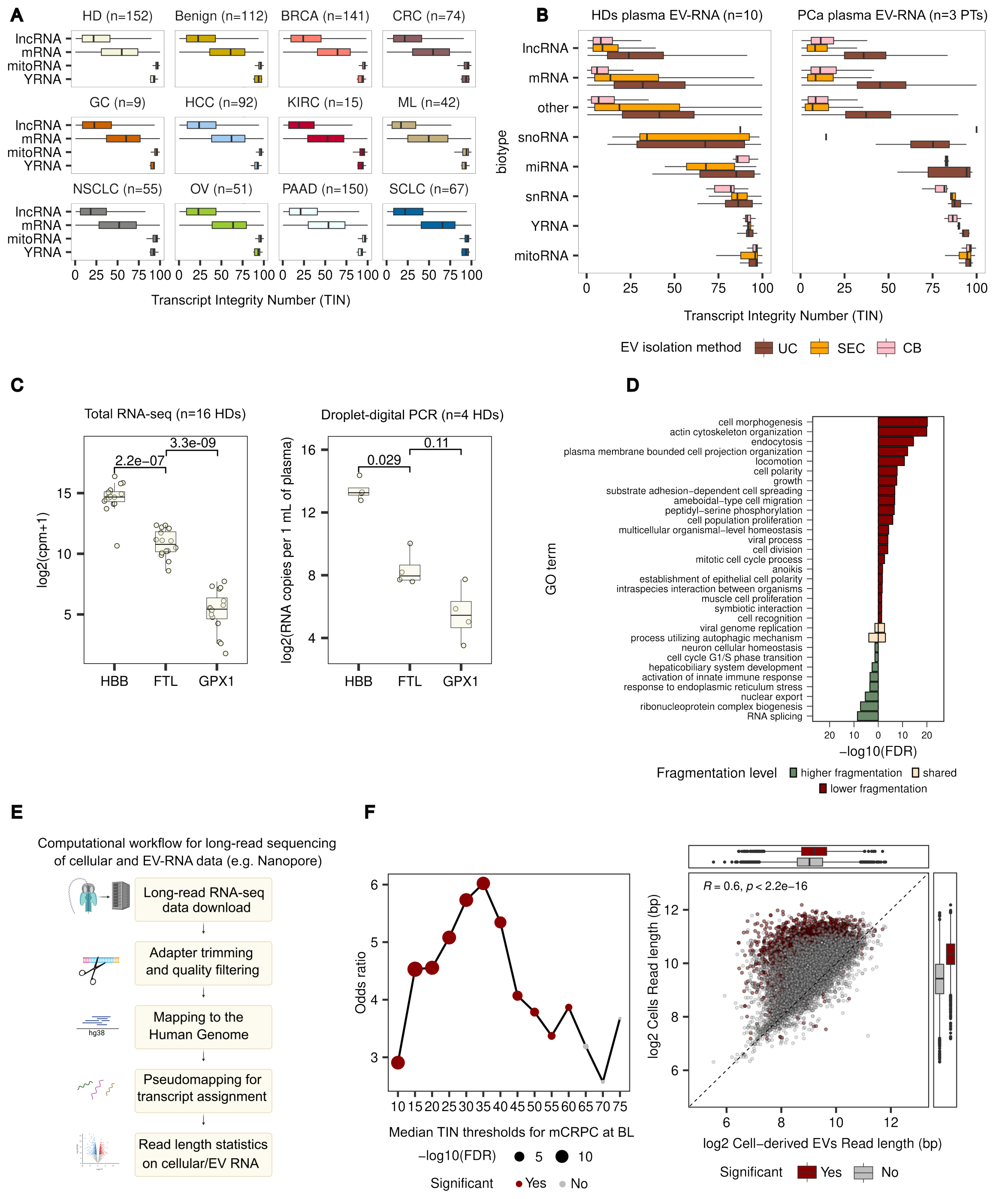


**Supplementary Figure 2.1: Validation of transcripts integrity**

A) Boxplots of TIN values of plasma EV-RNA from HDs and different tumor types from the independent Zhao cohort (total of 960 individuals). B) Boxplots of TIN values of n=10 HDs (left) and n=3 PCa patients (right) EV-RNA upon three isolation methods (UC=Ultracentrifugation, SEC= Size Exclusion Chromatography, CB=Charged-Based). C) Boxplots of HBB, FTL and GPX1 by total RNA-seq across n=16 study HDs (left) and by droplet-digital PCR (ddPCR) in n=4 HDs (right). Wilcoxon's two-sided test is reported. D) GO analysis based on differential TIN levels (n=54 mCRPC at BL, FDR < 0.05). E) Computational workflow for long-read RNA-seq data. F) Left, line plot showing Fisher's exact test for different median TIN thresholds for patients at BL. Each threshold stratified the transcripts into high and low TIN and in turn into closer to zero ($-0.5 < \log2FC < 0.5$) and far from zero ($\log2FC \leq 0.5$ or $\log2FC \geq 0.5$). The resulting contingency table has been used to run the test. Significant and non-significant data points are colored in red and grey, respectively. The dot size is proportional to the -log10(FDR) significance. Right, scatterplot showing cells (y axis) and cell-derived EVs (x axis) read lengths. Marginal boxplots show significant and non-significant data points, in red and grey, respectively. Abbreviations: TIN, Transcript Integrity Number; EV, extracellular vesicle; HD, healthy donor; PCa, prostate cancer; UC, ultracentrifugation; SEC, size-exclusion chromatography; CB, charge-based; HBB, hemoglobin subunit beta; FTL, ferritin light chain; GPX1, glutathione peroxidase 1; ddPCR, droplet digital PCR; GO, Gene Ontology; mCRPC, metastatic castration-resistant prostate cancer; BL, baseline; FDR, false discovery rate; lo2FC, log2FC fold change.

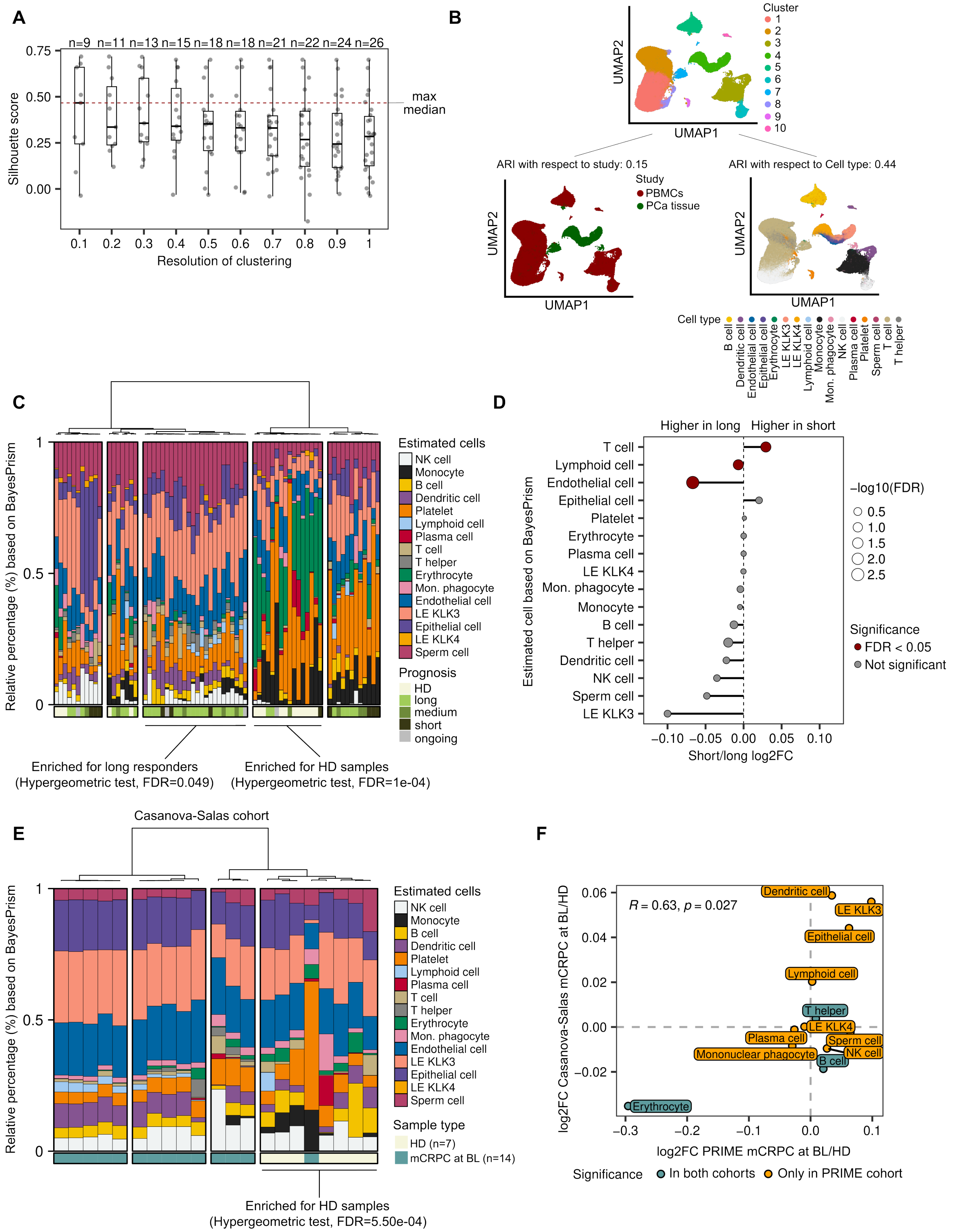


**Supplementary Figure 3.1: Plasma EV-RNA transcriptome deconvolution by BayesPrism**
A) Clustering based on random subsets of n=5,000 cells from scRNAseq reference based on the shared nearest neighbor (SNN) algorithm at different values of resolution on the x-axis. For

each cluster, a Silhouette score is computed (y-axis). The maximum median Silhouette score is reported as a red dotted horizontal line and number of clusters at each resolution values is reported on top. B) Umap showing the clustering with the optimal resolution parameter, colored by cluster (on top). The goodness of clustering was evaluated with respect to assay (bottom left) and cell type (bottom right) by calculating the Adjusted Rand Index (ARI). The ARI of clusters with respect to cell type (0.44) is higher than that of the assay (0.15), suggesting a negligible batch effect related to the assay. C) Clustering of BayesPrism deconvolution on PRIME enza cohort based on Aitchison distance. The optimal number of clusters was determined using a dynamic tree-cutting approach (a hybrid method). Significance based on the Hypergeometric test is reported for long responders and HDs in concordance with the third and fourth clusters, respectively. D) Lollipop plot comparing BayesPrism populations between short and long responders. The x-axis shows the log2FC between the two groups. Dot colors represent statistical significance, while dot size is proportional to -log10 of the FDR. E) Clustering of BayesPrism deconvolution on Casanova-Salas cohort based on Aitchison distance. The optimal number of clusters was determined using a dynamic tree-cutting approach (a hybrid method). Significance for Hypergeometric test is reported for HDs in concordance with the most-right cluster. F) Scatterplot showing the log2FC between mCRPC at BL and HDs for the PRIME Enza (x axis) and Casanova-Salas cohort (y axis). Deconvolution populations are colored by significance in both cohorts (blue) and in PRIME cohort only (orange). Spearman's correlation test statistic and p-value are reported. Abbreviations: EV, extracellular vesicle; scRNA-seq, single-cell RNA sequencing; SNN, shared nearest neighbor; UMAP, Uniform Manifold Approximation and Projection; ARI, Adjusted Rand Index; Enza, enzalutamide; HDs, healthy donors; $\log_2$FC, $\log_2$ fold change; FDR, false discovery rate; mCRPC, metastatic castration-resistant prostate cancer; BL, baseline.

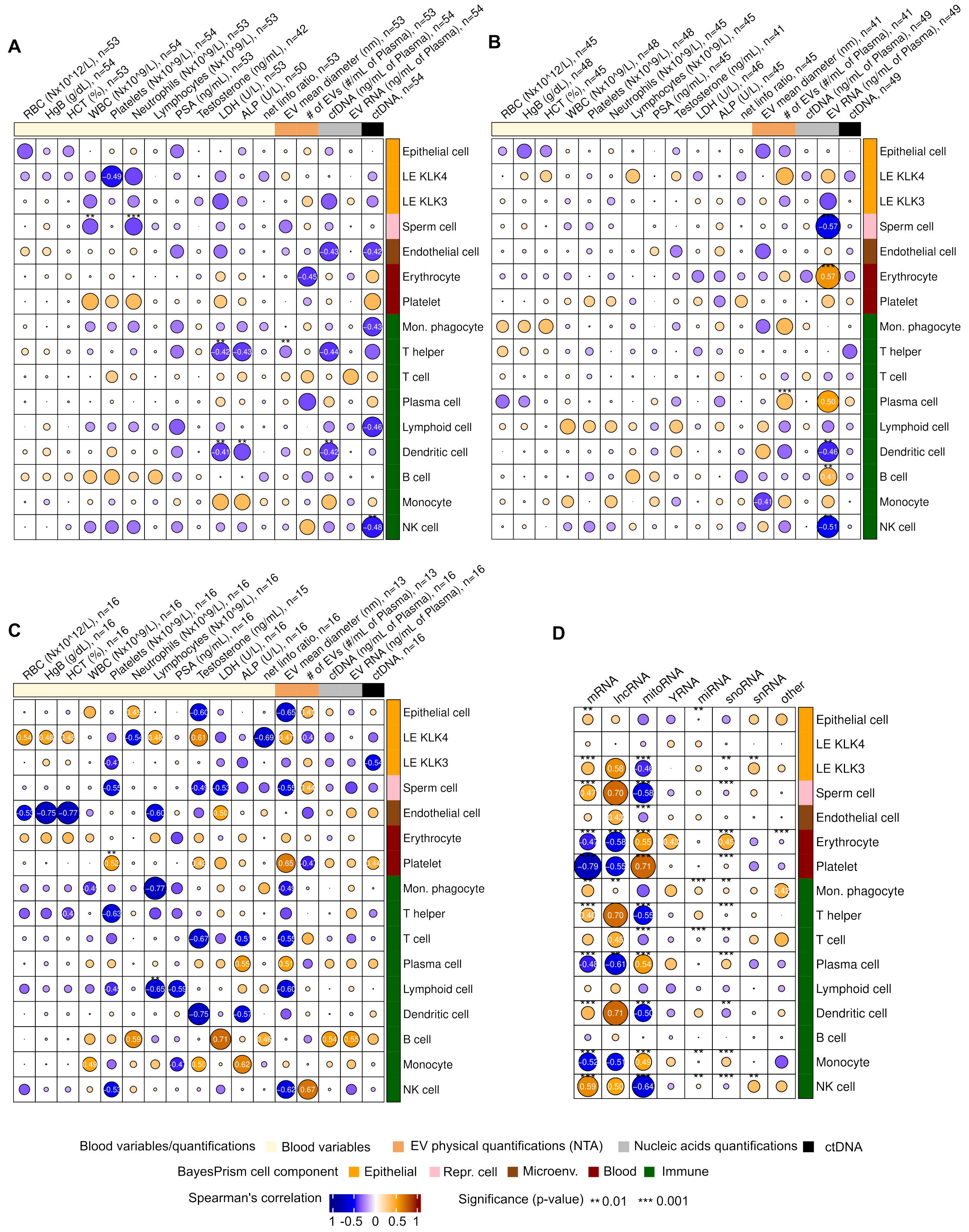


**Supplementary Figure 3.2: Correlations of BayesPrism deconvolution output with blood counts and biotypes**

Correlation matrix of blood counts with respect to BayesPrism populations at BL (A), 12w (B) and PR (C). Right annotation shows single cell categories, while the top one shows the blood counts and quantifications. Asterisks shows p-value significance for Spearman' correlation test at 0.01 (**) and

0.001 (***). B) Correlation matrix of blood counts with respect to BayesPrism populations at 12w. Right annotation shows single cell categories, while the top one shows the blood counts and quantifications. Asterisks p-value significance for Spearman's correlation test at 0.01 (**) and 0.001 (***). C) Correlation matrix of blood counts with respect to BayesPrism populations at PR. Right annotation shows single cell categories, while the top one shows the blood counts and quantifications. Asterisks show p-value significance for Spearman's correlation test at 0.01 (**) and 0.001 (***). D) Correlation matrix of BayesPrism output with respect to % of cpm per biotype. The annotation to the right shows single-cell categories. Asterisks show p-value significance for Spearman's correlation test at 0.01 (**) and 0.001 (***). Abbreviations: BL, baseline; 12w, 12 weeks; PR, progression; cpm, counts per million.

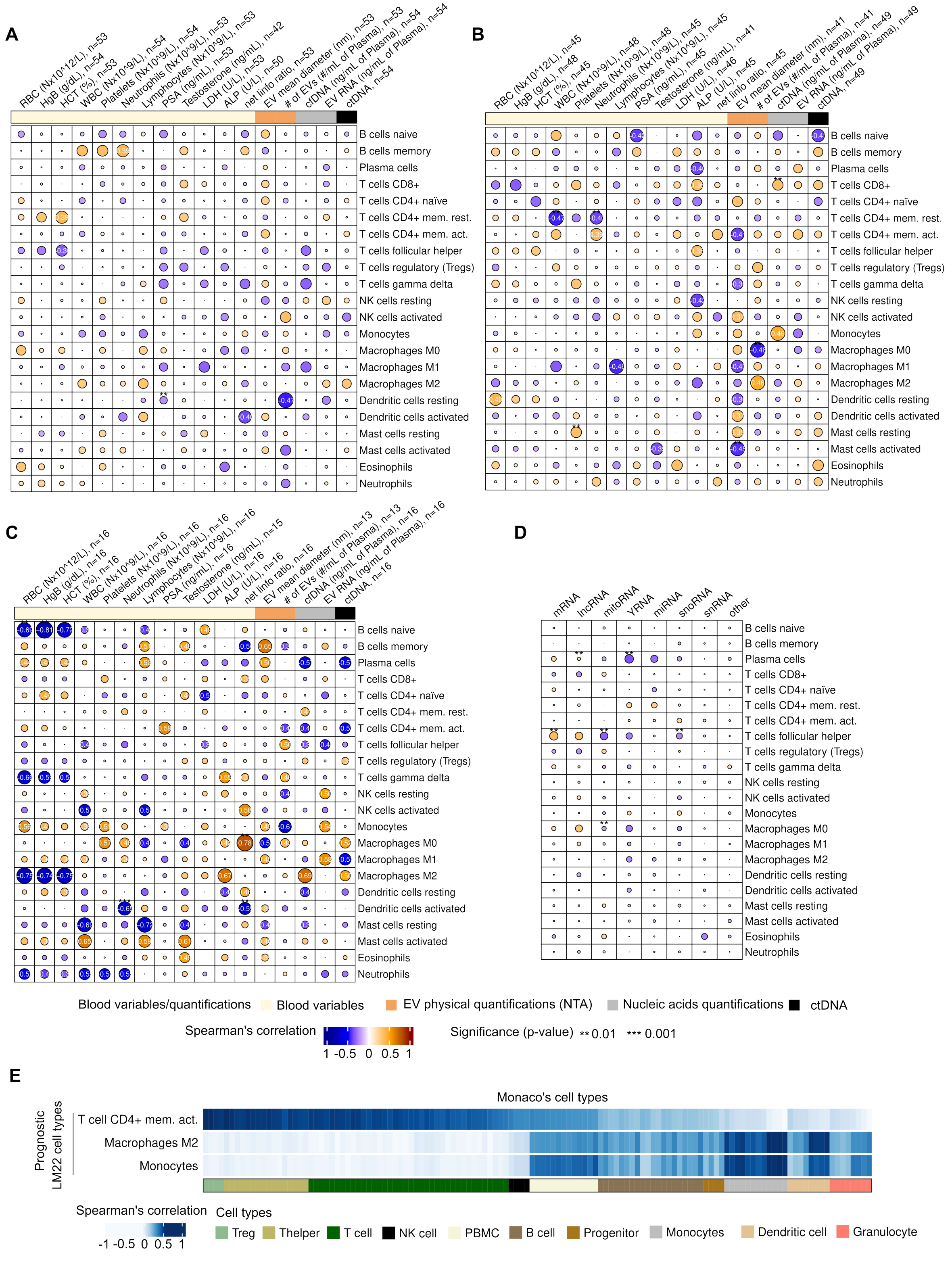


**Supplementary Figure 3.3: Correlations of CIBERSORT deconvolution output with blood counts and biotypes**

A) Correlation matrix of blood counts with respect to CIBERSORT populations at BL. Top annotation shows blood counts and quantifications. Asterisks show p-value significance for Spearman’s correlation test at 0.01 (**) and 0.001 (***). B) Correlation matrix of blood counts with respect to CIBERSORT populations at 12w. Top annotation shows blood counts and quantifications. Asterisks show p-value significance for Spearman’s correlation test at 0.01 (**) and 0.001 (***). C) Correlation matrix of blood counts with respect to CIBERSORT populations at PR. Top annotation shows blood counts and quantifications. Asterisks show p-value significance for Spearman’s correlation test at 0.01 (**) and 0.001 (***). D) Correlation matrix of CIBERSORT output with respect to % of cpm per biotype. Asterisks show p-value significance for Spearman’s correlation test at 0.01 (**) and 0.001 (***). E) Heatmap showing Spearman's correlations between immune populations identified in the PRIME mCRPC cohort (y-axis) and Monaco's samples (x-axis). The heatmap is annotated for Monaco's cell types at reduced granularity and sorted by the median correlation between CD4+ memory activated T cells and Monaco's cell types. Abbreviations: BL, baseline; 12w, 12 weeks; PR, progression; CPM, counts per million; mCRPC, metastatic castration-resistant prostate cancer.

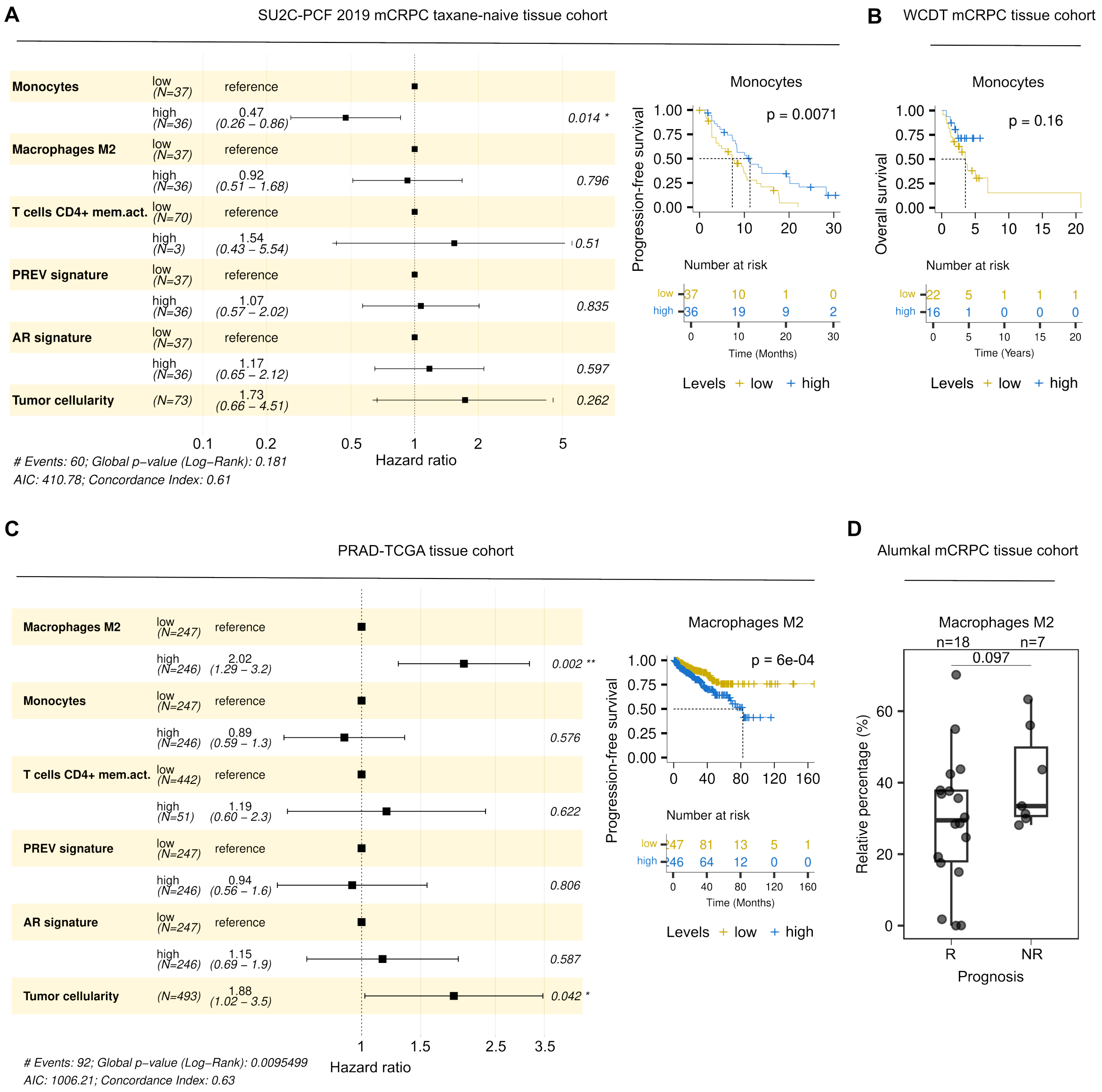


**Supplementary Figure 3.4: The prognostic value of plasma EV-related signal in mCRPC tissue**

A) Multivariate Cox proportional Hazard ratio analysis in SU2C-PCF 2019 mCRPC taxane-naïve tissue cohort (n=73, left). PFS curve of Monoctyes estimated with CIBERSORT in SU2C-PCF 2019 mCRPC taxane-naïve tissue cohort (right). B) Kaplan-Meier curves showing overall survival (OS) according to Monocytes levels in ARSI-naïve patients from Weast Coast Dream Team (WCDT) cohrort treated with Enzalutamide (n=38 pts). C) Multivariate Cox proportional Hazard ratio analysis in PRAD-TCGA PCa tissue cohort (n=493, left). PFS curve of Macrophages M2 estimated with CIBERSORT in PRAD-TCGA cohort (right). D) Boxplots showing Macrophages M2 levels estimated with CIBERSORT in the Alumkal mCRPC tissue cohort , stratified by prognosis status (R, responders; NR, not-responders). Wilcoxon Mann-Whitney two-sided p-value is reported. Abbreviations: EV, extracellular vesicle; mCRPC, metastatic castration-resistant prostate cancer; SU2C-PCF, Stand Up To Cancer–Prostate Cancer Foundation; PFS, progression-free survival; OS, overall survival; ARSI, androgen receptor signaling inhibitor; WCDT, West Coast Dream Team; PRAD, prostate adenocarcinoma; TCGA, The Cancer Genome Atlas; PCa, prostate cancer; R, responders; NR, non-responders.

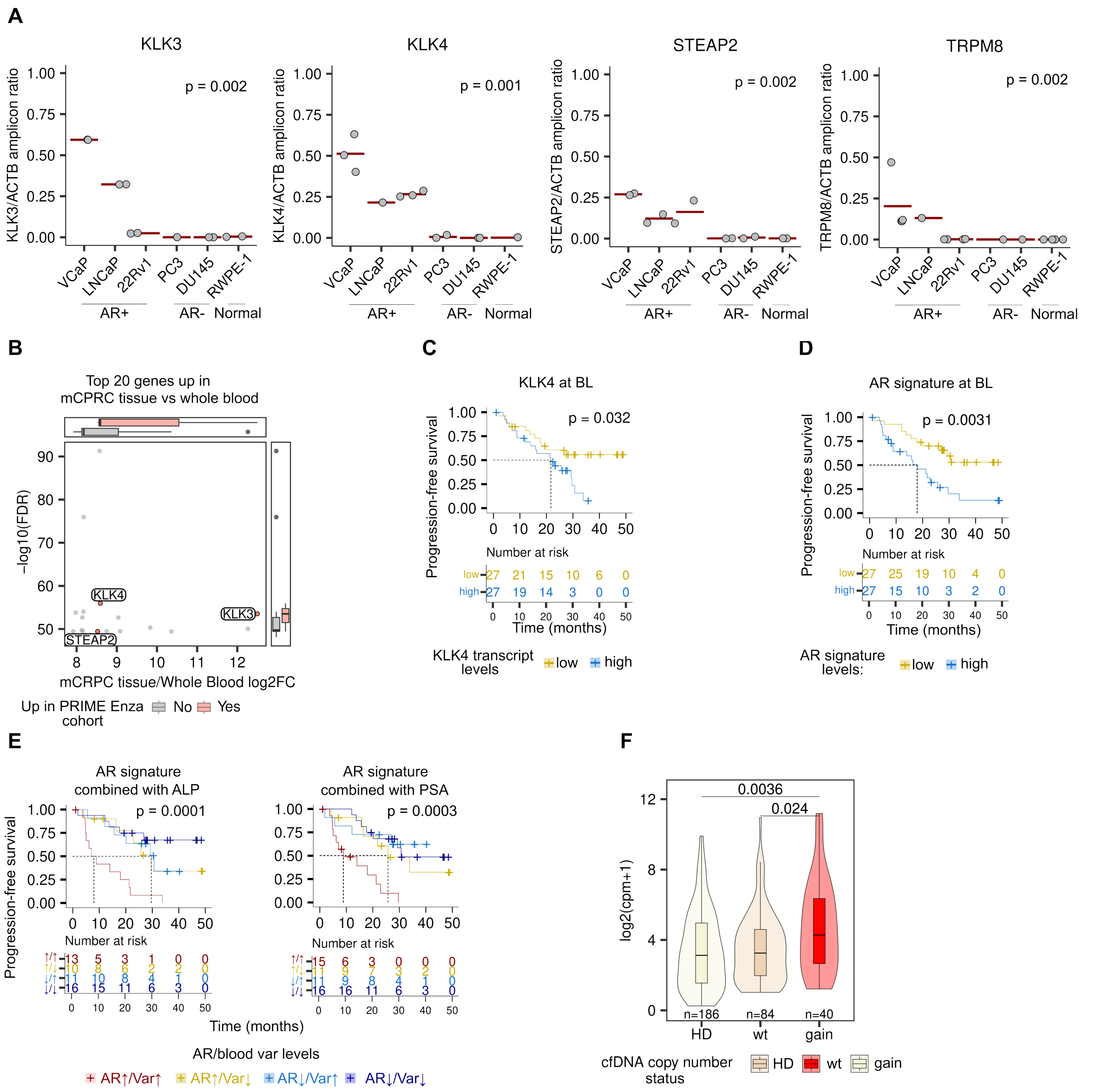


**Supplementary Figure 4.1: Plasma EVs carry a tumor-related signal**
A) Levels of PCa-related transcripts by droplet-digital PCR (ddPCR) in PCa cell-derived EVs. The cDNA abundance is represented as the ratio with respect to ACTB (y-axis) for cell-derived EVs grouped by AR status (x-axis); red bars indicate mean values; Wilcoxon's Mann-Whitney test is reported comparing AR+ and AR- cell-derived EVs. Biological replicates: KLK3, n=2; KLK4, n=4, except LNCaP (n=1) and DU145 (n=2); STEAP, n=2; TRPM8, n=4, except LNCaP, PC3, and DU145 (n=1). B) Scatterplot showing top 20 genes upregulated in mCRPC tissue (n=88 samples) with respect to whole blood (n=559 samples). On the y-axis, the significance of the analyses is reported as -log10 of FDR, on the x-axis, the log2 fold change. Dots are colored by significance for comparison

between PRIME Enza cohort at BL vs HDs plasma EV-RNA. C) Kaplan-Meier curves of KLK4 transcript levels in PRIME Enza mCPRC plasma EV-RNA at BL. Stratification based on median. D) Kaplan-Meier survival curves of the AR signature at BL in PRIME Enza cohort plasma EV-RNA at BL. E) Kaplan-Meier curves of AR signature combined with ALP (left) and PSA (right) in PRIME Enza cohort at BL. Only curves with p-value < 0.01 are shown. Stratification of combinations based on median levels of AR signature and blood variables. F) EV transcript levels of oncogenes in mCPRC at BL and HD. mCPRC data is grouped based on the underlying genomic status of each gene as the matched cfDNA profiling; Wilcoxon's Mann-Whitney two-sided test is reported on top. Abbreviations: PCa, prostate cancer; ddPCR, droplet digital PCR; cDNA, complementary DNA; EV, extracellular vesicle; AR, androgen receptor; mCRPC, metastatic castration-resistant prostate cancer; FDR, false discovery rate; log2FC, log2FC fold change; BL, baseline; HD, healthy donor; cfDNA, cell-free DNA; ALP, alkaline phosphatase; PSA, prostate-specific antigen.

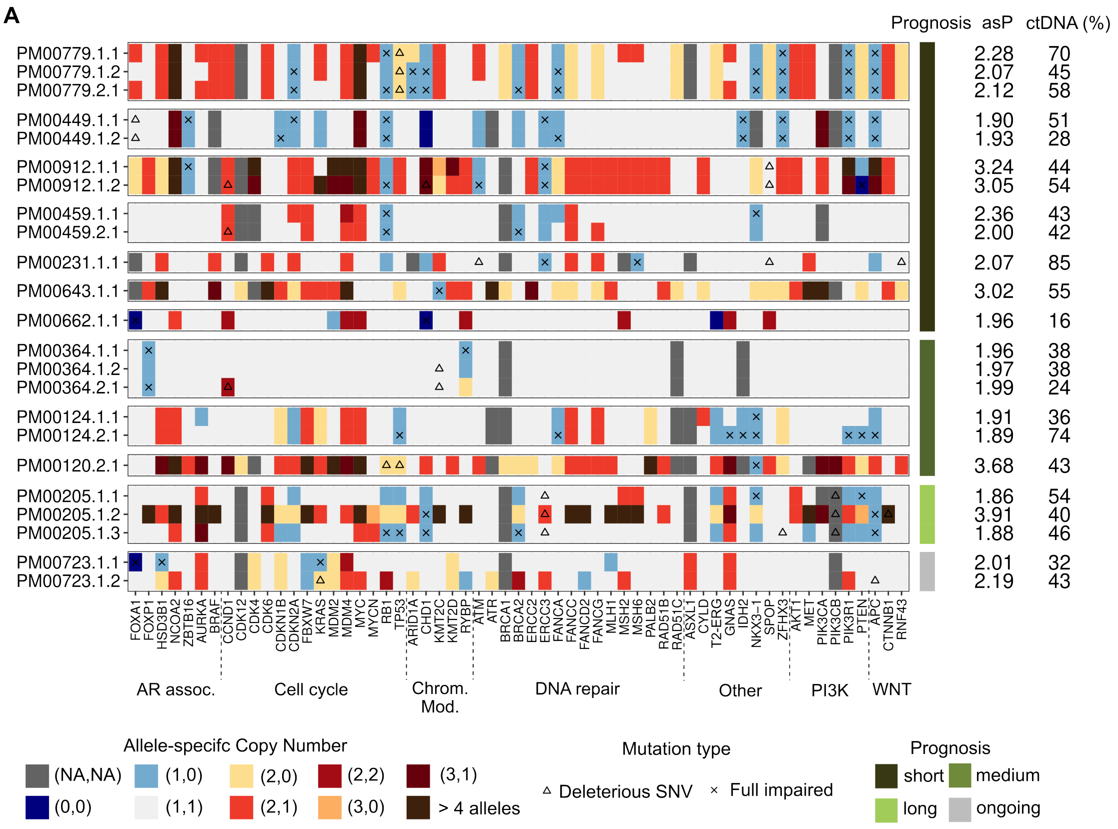


**Supplementary Figure 4.2: Oncoprint of allele-specific copy number calls from plasma ctDNA**

A) Oncoprint showing allele-specific copy number calls for PRIME Enza mCRPC patients with ctDNA > 15%. Detectable ctDNA at BL (top, n=15), 12w (middle, n=6) and at PR (bottom, n=10). The plot is annotated from prognosis (left bar), ctDNA and tumor Ploidy levels (right bar) and oncogene/tumor suppressor annotation (bottom bar). Abbreviations: ctDNA, circulating tumor DNA; mCRPC, metastatic castration-resistant prostate cancer; BL, baseline; 12w, 12 weeks; PR, progression.

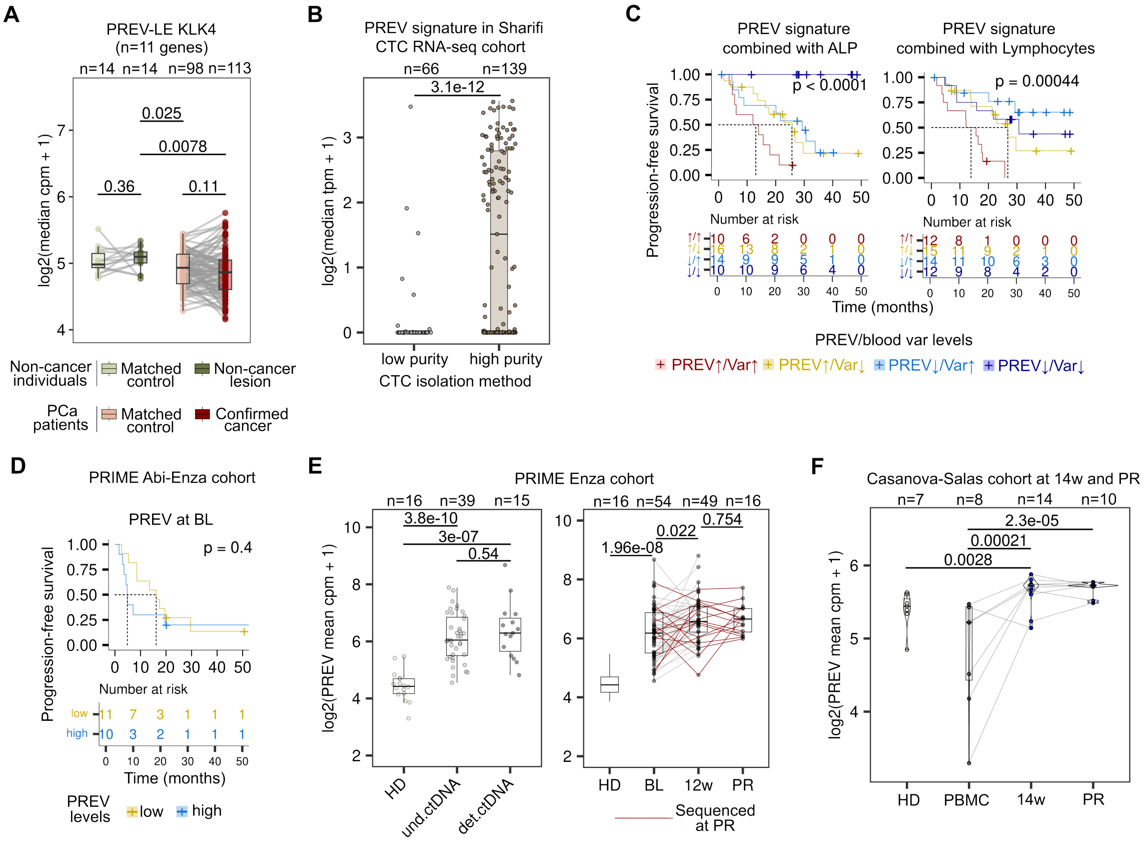


**Supplementary Figure 5.1: The characterization of PREV signature across blood components and advanced PCa cohorts:** A) PREV expression of cell-type specific PREV genes in the ReIMAGINE cohort, selected using the Tuong dataset, including PREV genes highly expressed in LE KLK4 (n=11); Wilcoxon's Mann-Whitney two-sided paired test is reported for paired individuals, while Mann-Whitney two-sided unpaired test is reported for unpaired biopsies. B) Boxplots showing PREV signature levels in low versus high purity circulating tumor cells (CTCs) from the Sharifi RNAseq cohort of advanced PCa (n=205). C) Kaplan-Meier curves of PREV signature combined with ALP (left) and Lymphocyte counts (right) in PRIME Enza cohort at BL. Only curves with p-value < 0.01 are shown. Stratification of combinations based on median levels of PREV signature and blood variables. D) Kaplan-Meier curves of PREV levels plasma EV-RNA of the PRIME Abi-Enza cohort at BL. Stratification based on median. E) Left, boxplots showing the PREV levels in PRIME Enza cohort plasma EV-RNA at BL stratified by ctDNA positivity, and HDs. Wilcoxon's two-sided test is reported. Right, PREV levels at different time points and HDs. Two-sided paired Wilcoxon's test was used for comparisons across time points; unpaired Wilcoxon's test was used for HDs vs patients. F) PREV signature levels on Casanova-Salas cohort at 14 weeks (14w) and disease progression (PR) compared with HD and PBMCs. One-side paired Wilcoxon's test was used for comparisons across time points; unpaired Wilcoxon's test was used for HDs vs patients. Abbreviations: PCa, prostate cancer; CTCs, circulating tumor cells; ALP, alkaline phosphatase; BL, baseline; EV-RNA, extracellular vesicle RNA; Abi, abiraterone; Enza, enzalutamide; ctDNA, circulating tumor DNA; HD, healthy donor; 14w, 14 weeks; PR, progression; PBMCs, peripheral blood mononuclear cells.

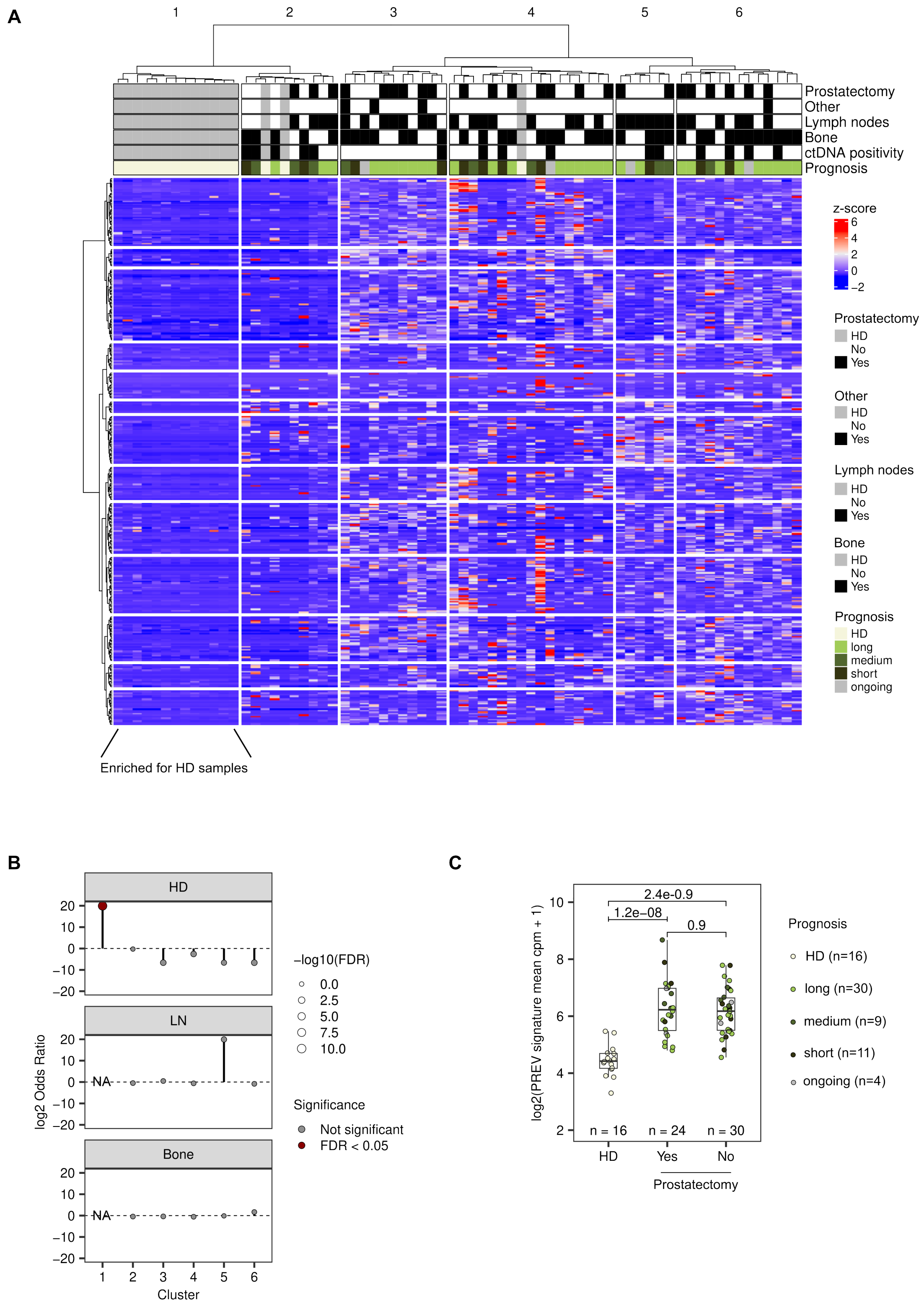


**Supplementary Figure 5.2: The PREV stratifies patients from HDs and is independent of prostatectomy status in the PRIME Enza cohort:** A) Heatmap and dendrogram of the PRIME Enza cohort patients at BL (n=54) and HDs (n=16) based on the z-score cpm values of the

PREV signature (n=353 genes). For the clustering, Spearman correlation distance and Ward's linkage method were used. The optimal number of clusters was determined using a dynamic tree-cutting approach (a hybrid method). Sample/patient annotations include prostatectomy, site of metastasis, and prognosis (top bars). B) Lollipop plot showing enrichment for each cluster by Fisher's test in panel A (i.e., whether a cluster is significantly enriched for HDs, lymph nodes, or bone metastases). On y axis is reported the log2 Odds ratio, on x the cluster. Dots color shows FDR significance. C) PREV signature levels in the PRIME Enza cohort, stratified by prostatectomy status (Yes/No), compared to HDs. Wilcoxon Mann Withney two-sided p-value is reported. Abbreviations: HD, healthy donor; Enza, enzalutamide; BL, baseline; cpm, count per million; FDR, false discovery rate.

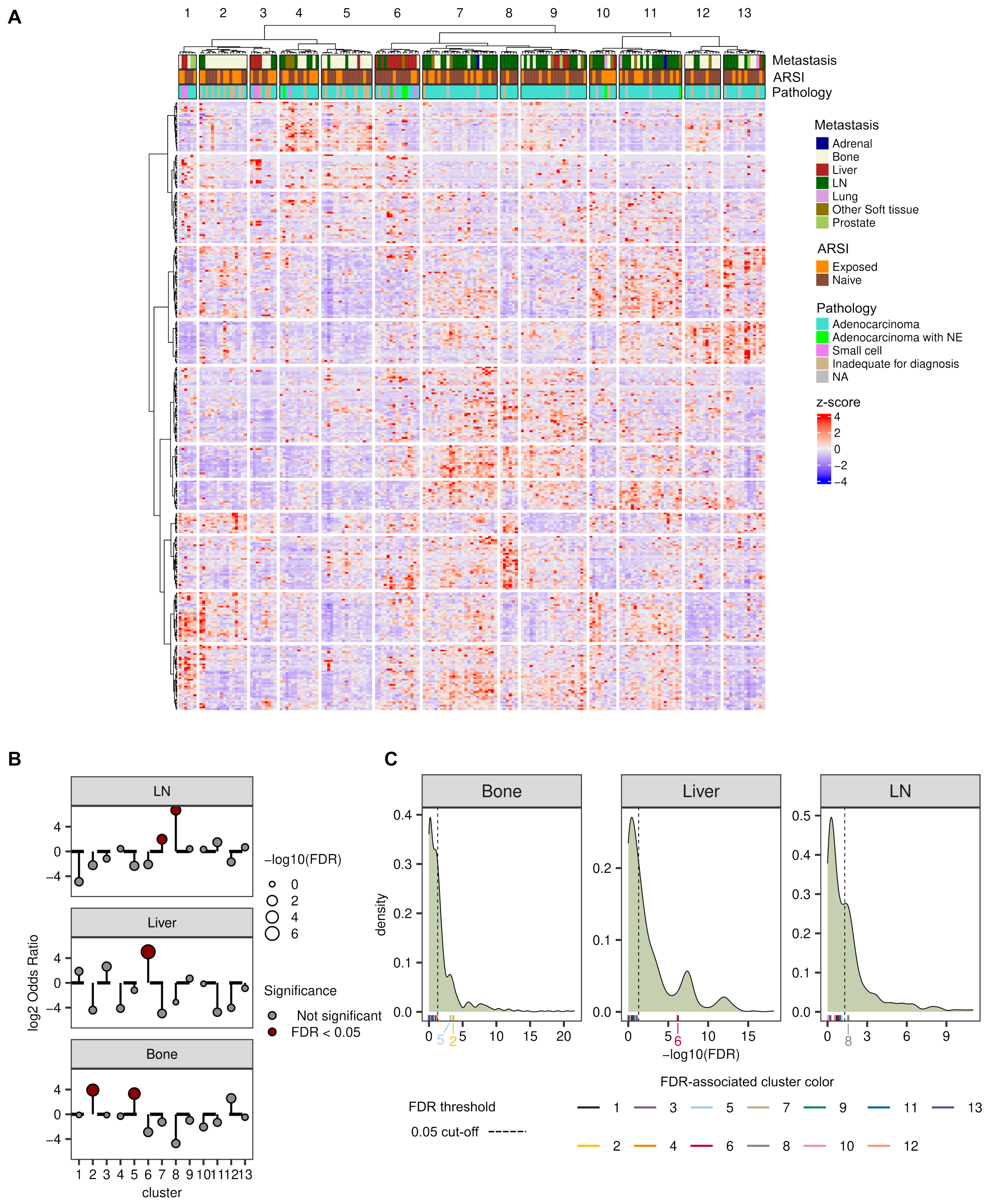


**Supplementary Figure 5.3: The PREV stratifies the site of metastasis in SU2C-PCF 2019 taxane naïve cohort:** A) Heatmap showing the clustering of patients based on z-score fpkm values of the PREV signature (n=336 genes) in SU2C-PCF 2019 mCRPC tissue cohort. The plot annotated for site of metastasis, ARSI exposure and pathology (top bars). For the clustering, Spearman's correlation distance and Ward's linkage method were used. The optimal number of clusters was determined using a dynamic tree cutting approach (hybrid method). B) Lollipop plot showing enrichment for each cluster by Fisher's test in panel A. On y axis is reported the log2 Odds ratio, on x the cluster. Each facet is showing each of the site of metastasis tested. Dots color shows FDR significance. C) Density plot representing the -log10 FDR distribution of Fisher's test for randomly selected genes (n=363 genes, 100 times) from a set of genes upregulated in mCRPC tissue with respect to whole blood (n=10,002 genes). Each facet represents a site of metastasis tested; lower vertical lines are positioned according to -log10 FDR and colored by cluster reported in panel B for PREV signature. Vertical dotted intercepts represent 0.05 FDR cut-offs. Abbreviations: SU2C-PCF, Stand Up To Cancer–Prostate Cancer Foundation; mCRPC, metastatic castration-resistant prostate cancer; FPKM, fragment per kilobase of transcript per million mapped read; ARSI, androgen receptor signaling inhibitor; FDR, false discovery rate.
